# Metabolic Reprogramming via targeting ACOD1 promotes polarization and anti-tumor activity of human CAR-iMACs in solid tumors

**DOI:** 10.1101/2023.04.20.537647

**Authors:** Xudong Wang, Siyu Su, Yuqing Zhu, Xiaolong Cheng, Chen Cheng, Leilei Chen, Anhua Lei, Li Zhang, Yuyan Xu, Wei Li, Yi Zhang, Dan Ye, Jin Zhang

**Affiliations:** Center for Stem Cell and Regenerative Medicine, Department of Basic Medical Sciences, and Bone Marrow for Transplantation Center of the First Affiliated Hospital; Zhejiang University School of Medicine, Hangzhou, 310058, China; Zhejiang Laboratory for Systems & Precision Medicine, Zhejiang University Medical Center; Hangzhou, Zhejiang Province, 311121, China; Quanzhou First Hospital Affiliated to Fujian Medical University, Quanzhou 362000, China; Center for Stem Cell and Translational Medicine, School of Life Sciences, Anhui University, Hefei, Anhui 230601, P.R. China; Center for Genetic Medicine Research, Children’s National Hospital, 111 Michigan Ave NW, Washington, DC, 20010, USA; Department of Genomics and Precision Medicine, George Washington University, 111 Michigan Ave NW, Washington, DC, 20010, USA; Shanghai Key Laboratory of Clinical Geriatric Medicine, Shanghai, Huadong Hospital, and Shanghai Key laboratory of Medical Epigenetics, International Co-laboratory of Medical Epigenetics and Metabolism (Ministry of Science and Technology), and Molecular and Cell Biology Lab, Institutes of Biomedical Sciences, Shanghai Medical College of Fudan University, Shanghai, 200032, China; CellOrigin Inc., Hangzhou, China; Department of General Surgery, Huashan Hospital, Fudan University, Shanghai 200040, China

## Abstract

The pro-inflammatory state of macrophages is crucial in conferring its role in combating tumor cells. That state is closely associated with metabolic reprogramming. Here we identified key metabolic genes regulating macrophage pro-inflammatory activation in a pooled metabolic gene knockout CRISPR screen. We found that *KEAP1 a*nd *ACOD1* are strong regulators of the pro-inflammatory state, and therefore developed human *ACOD1* knockout macrophages with our induced pluripotent stem cell-derived CAR-macrophage (CAR-iMAC) platform. The engineered iMACs showed stronger and more persistent polarization toward the pro-inflammatory state, more ROS production, and more potent phagocytosis and cytotoxic functions against cancer cells *in vitro*. Upon transplantation to ovarian or pancreatic cancer mouse models, *ACOD1* depleted CAR-iMACs exhibited enhanced capacity in repressing tumors *in vivo* and prolonged the lifespan of mice. In addition, combining *ACOD1-*depleted CAR-iMACs with immune check point inhibitors (ICIs), such as the anti-CD47 antibody or anti-PD1 antibody resulted in stronger tumor suppressing effect. Mechanistically, the depletion of ACOD1 reduced the immunometabolite itaconate, allowing KEAP1 to prevent NRF2 from entering the nucleus to activate the anti-inflammatory program. This study demonstrates that ACOD1 is a new myeloid target for cancer immunotherapy and metabolically engineered human iPSC-derived CAR-iMACs exhibit enhanced polarization and anti-tumor functions in adoptive cell transfer therapies.

## Introduction

Macrophages serve as the first line of host defense and play a key role in innate immunity. The primary function of macrophages is phagocytosis and microbial killing^1^. They also participate in a variety of physiological and pathological processes such as development, inflammation and tumorigenesis. Macrophages can be generally defined into two highly plastic states: LPS and IFN-γ-activated pro-inflammatory macrophages (M1-like macrophages) and IL-4 or IL-10 induced alternatively activated macrophages (M2-like macrophages)^2^. Recent studies revealed different metabolic pathways are closely associated with the different states. Pro-inflammatory macrophages mainly rely on glycolysis, exhibit the impaired tricarboxylic acid (TCA) cycle and express the Inducible Nitric Oxide Synthase (iNOS), whereas alternatively activated macrophages mainly rely on mitochondrial oxidative phosphorylation (OXPHOS)^3^.

Macrophages are highly plastic cells that can adapt to their surrounding environment. Pro-inflammatory M1-like macrophages play a crucial role in responding to viruses and bacteria infection and participate in anti-tumor immunity^4^, whereas M2-like macrophages can contribute to tumor progression^4^. Tumors can recruit and reprogram macrophages to become the M2-like tumor-associated macrophages (TAMs). TAMs suppress endogenous cytotoxic T cells, secrete chemokines to recruit Treg cells^5^, and secrete factors such as VEGF and matrix metalloproteinase enzymes to remodel the TME, promoting tumor angiogenesis and metastasis^6^. Thus, a primary goal of macrophage-based cancer immunotherapy is to reduce anti-inflammatory macrophages and increase pro-inflammatory macrophages.

One of the strategies targeting macrophages is to inhibit TAMs *in situ* in the TME. For instance, an inhibitor of the CSF-1 receptor (CSF-1R) could significantly reduce TAMs and block glioma progression in a mouse model^7^. An alternative strategy is to modify macrophages *ex vivo* through genetically engineered monocytes and macrophages, and the engineered macrophages can be adoptively transferred to tumor-carrying mice. A modified lentiviral vector, Vpx-LV^8^, and chimeric adenoviral vector Ad5f35^9^ were used to efficiently transduce primary monocytes and macrophages. We developed the iPSC-derived engineered CAR-macrophage (CAR-iMAC), which may become a powerful source of engineered macrophage for immunotherapy due to its ease of engineering and adequate supply. We also demonstrated antigen-dependent anti-tumor functions when challenged with antigen-expressing cancer cells *in vitro* and *in vivo*^10^. However, the first generation of CAR-iMACs was not designed to assume a pro-inflammatory state, necessitating further engineering in this direction. In this study, we first used pooled CRISPR-Cas9 screens to identify the metabolic regulators of macrophage pro-inflammatory activation. Our screen revealed that the ACOD1/KEAP1/NRF2 pathway regulates cellular metabolism and pro-inflammatory activity of macrophages. Moreover, *ACOD1* depleted iMACs or CAR-iMACs are superior in comparison to the unmodified ones in cancer immunotherapies because of their enhanced *in vitro* and *in vivo* anti-tumor functions. Therefore, the present work highlights a new myeloid target in cancer immunotherapy and provides novel engineering strategies for adoptive cell transfer therapies using metabolically rewired CAR-macrophages.

## Results

### A CRISPR screen identified *KEAP1* deletion abrogated LPS and IFN-γ induced pro-inflammatory activation of macrophages

To identify the possible genes influencing macrophage pro-inflammatory activation, we designed a CRISPR screen^11^ using a human metabolic sgRNA library containing metabolism-related transcription factors, small molecule transporters, and metabolic enzymes in a Cas9-expressing lentiviral vector^12^. The THP-1 cell line is a convenient system for studying human macrophages *in vitro,* as the THP-1 cells could be induced into macrophages (tMACs) by PMA stimulation (Extended Data Fig. 1a) and could be further activated towards pro-inflammatory macrophages after LPS and IFN-γ stimulation (Extended Data Fig. 1b). No significant differences in the pro-inflammatory activation capacity were found between WT and the sgRNA library virus transduced THP-1 cells (Extended Data Fig. 1c). After transduction and selection, THP-1 cells were differentiated into macrophages and stimulated with LPS and IFN-γ for 24 h. CD80-high and CD80-low populations were sorted using flow cytometry. Top ranking candidate genes enriched in the two populations were unraveled using deep sequencing (Fig. 1a). GO analysis revealed that in the CD80-high population, sgRNAs/genes related to NAD activity, hypoxia, and amino acid transporter were enriched, whereas in the CD80-low population, sgRNAs/genes related to reactive oxygen species, glycosaminoglycan biosynthesis, and glycolysis were enriched (Extended Data Fig. 1d). The screen results were also visualized with a volcano plot, which revealed that the sgRNAs targeting *KEAP1* was significantly enriched in the CD80-low population (Fig. 1b), and sgRNA counts of *KEAP1* were significantly higher in the CD80-low population (Extended Data Fig. 1e). During our screen, sg*NFKB1*s were also enriched in the CD80-low population (Extended Data Fig. 1f left), which is consistent with its role in promoting the inflammation program^13^. When *NFKB1* was deleted in THP-1 cells (Extended Data Fig. 1g), the expression of CD80 was significantly abrogated in the LPS and IFN-γ-induced macrophages (Extended Data Fig. 1h,i). This demonstrated that the validity of our screen in THP-1 cells was credible. We subsequently deleted *KEAP1* in THP-1 cells to validate the effect of KEAP1 on macrophage pro-inflammatory activation. We designed three sgRNAs to target the human *KEAP1* (Extended Data Fig. 2a) with good efficiency (Extended Data Fig. 2b,c), and the KEAP1 level could be reduced in THP1 cells (Fig. 1c and Extended Data Fig. 2d). To examine activation of the *KEAP*1^-/-^ macrophages, we treated tMACs with LPS and IFN-γ for 2, 8, and 24 h. The expression of CD80 is significantly abrogated in *KEAP1*^-/-^ macrophages after 8 and 24 h of stimulation (Fig. 1d,e). The expression of pro-inflammatory genes was also reduced in *KEAP1*^-/-^ macrophages, with the maximal difference between WT and *KEAP1*^-/-^ macrophages observed after 8 h of stimulation (Fig. 1f). As sg*KEAP1*-3 showed the highest efficiency (Extended Data Fig. 2c,d), sg*KEAP1*-3 transduced tMACs were used for further analysis. With RNA-seq analysis, we identified genes related to Toll-like receptor signaling pathway, phagosome, NOD-like receptor signaling pathway, and NF-kappa B signaling pathway was higher in WT macrophages after stimulation (Extended Data Fig. 3a), whereas genes related to Glutathione metabolism and Oxidative phosphorylation was higher in knockout macrophages with the same simulation (Extended Data Fig. 3b). Together, these findings indicate that *KEAP1* deletion inhibits the pro-inflammatory activation of macrophages.

**Fig. 1.**
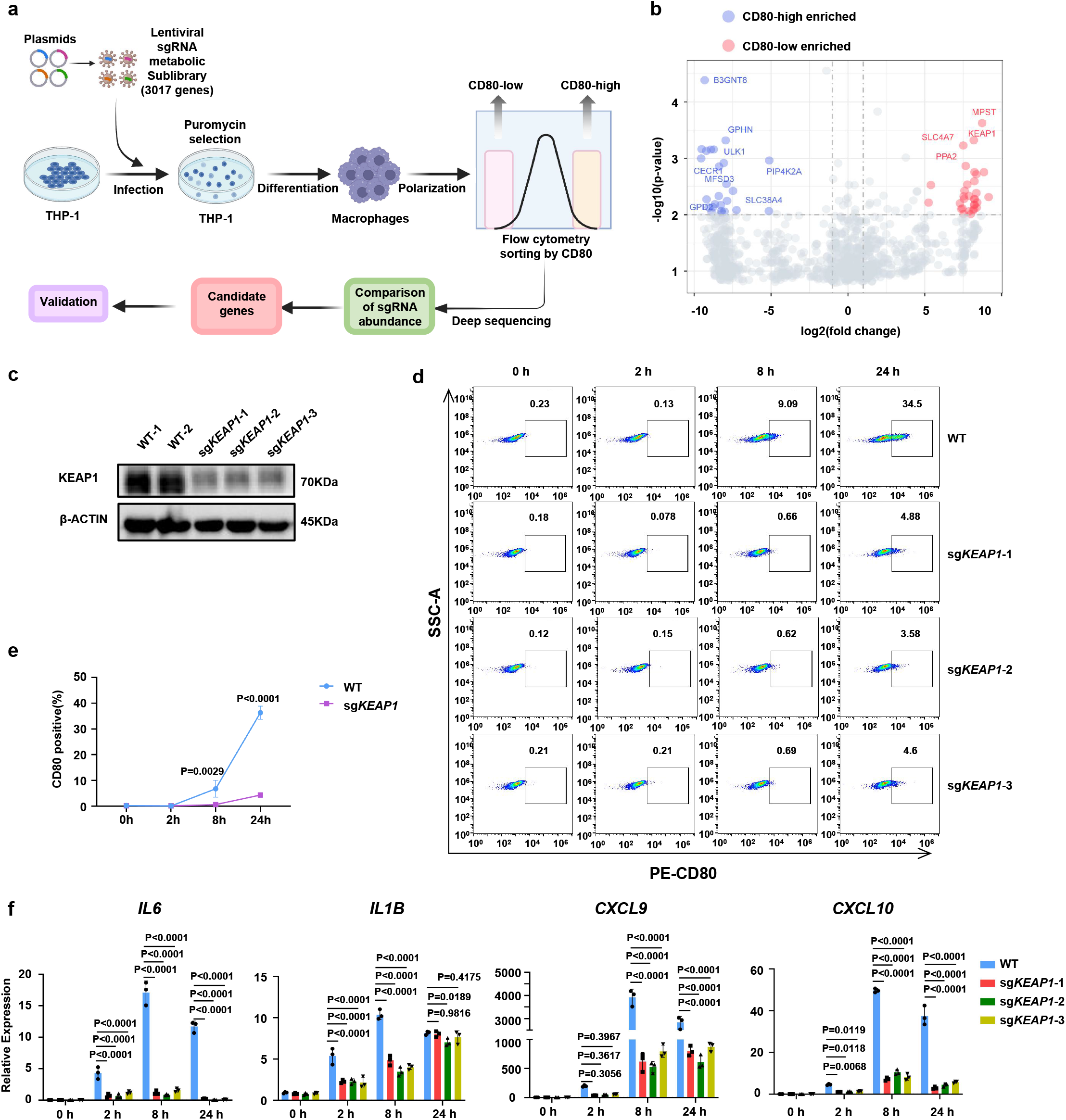
A CRISPR screen identified *KEAP1* deletion abrogated LPS and IFN-γ induced pro-inflammatory activation in macrophages. **a**, A schematic diagram of the pooled CRISPR screen of metabolic genes in THP1-induced macrophages or tMACs. **b**, A volcano plot displaying sgRNA-targeted genes enriched in the CD80-high (blue) and CD80-low (red) populations. **c**, The protein level of KEAP1 in WT and *KEAP1*-deficient tMACs. **d-e**, Flow cytometry plots and quantification of CD80 expression on WT and *KEAP1*-deficient tMACs after LPS and IFN-γ stimulation for 0, 2, 8, and 24 h. **f,** qRT-PCR analysis of IL6, IL1B, CXCL9, and CXCL10 expression in WT and *KEAP1*-deficient tMACs after LPS and IFN-γ stimulation at different time points (n=3). Data were repeated independently in three separate experiments. **e-f**, The data were displayed as mean ± SD. Statistics by two-way ANOVA test.

**Fig. 2.**
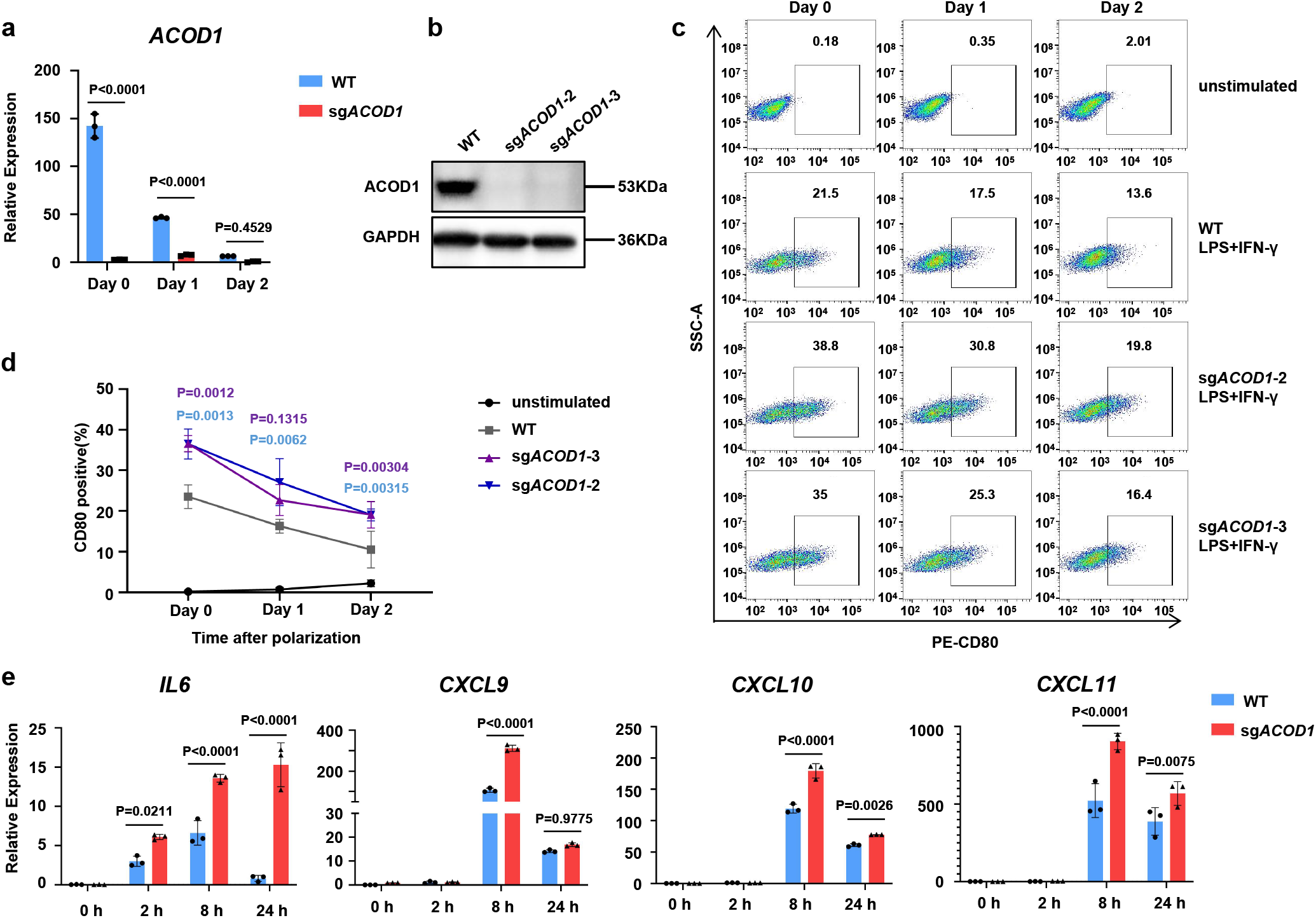
*ACOD1* deletion promoted pro-inflammatory activation in tMACs. **a,** The relative expression of *ACOD1* in WT and sgACOD1 transduced tMACs with LPS and IFN-γ stimulation at the indicated time points (n=3). **b**, The protein level of ACOD1 in WT and sgACOD1-transduced cells after LPS and IFN-γ stimulation for 24 h. **c,d**, Flow cytometry plots and quantification of CD80 expression in unstimulated, WT, and sg*ACOD1* transduced tMACs with indicated treatments (d, n=3). The tMACs were stimulated by 50 ng/mL LPS and 50 ng/mL IFN-γ for 24 h, then withdrawn from the stimulation and further cultured for 24 h (Day 1) or 48 h (Day 2). **e**, qRT-PCR for mRNA expression of pro-inflammatory genes in WT and sg*ACOD1* transduced tMACs after LPS and IFN-γ stimulation at different time points (n=3). **a, d, e**, Data was shown as mean ± SD. Statistics by two-way ANOVA test.

**Fig.3.**
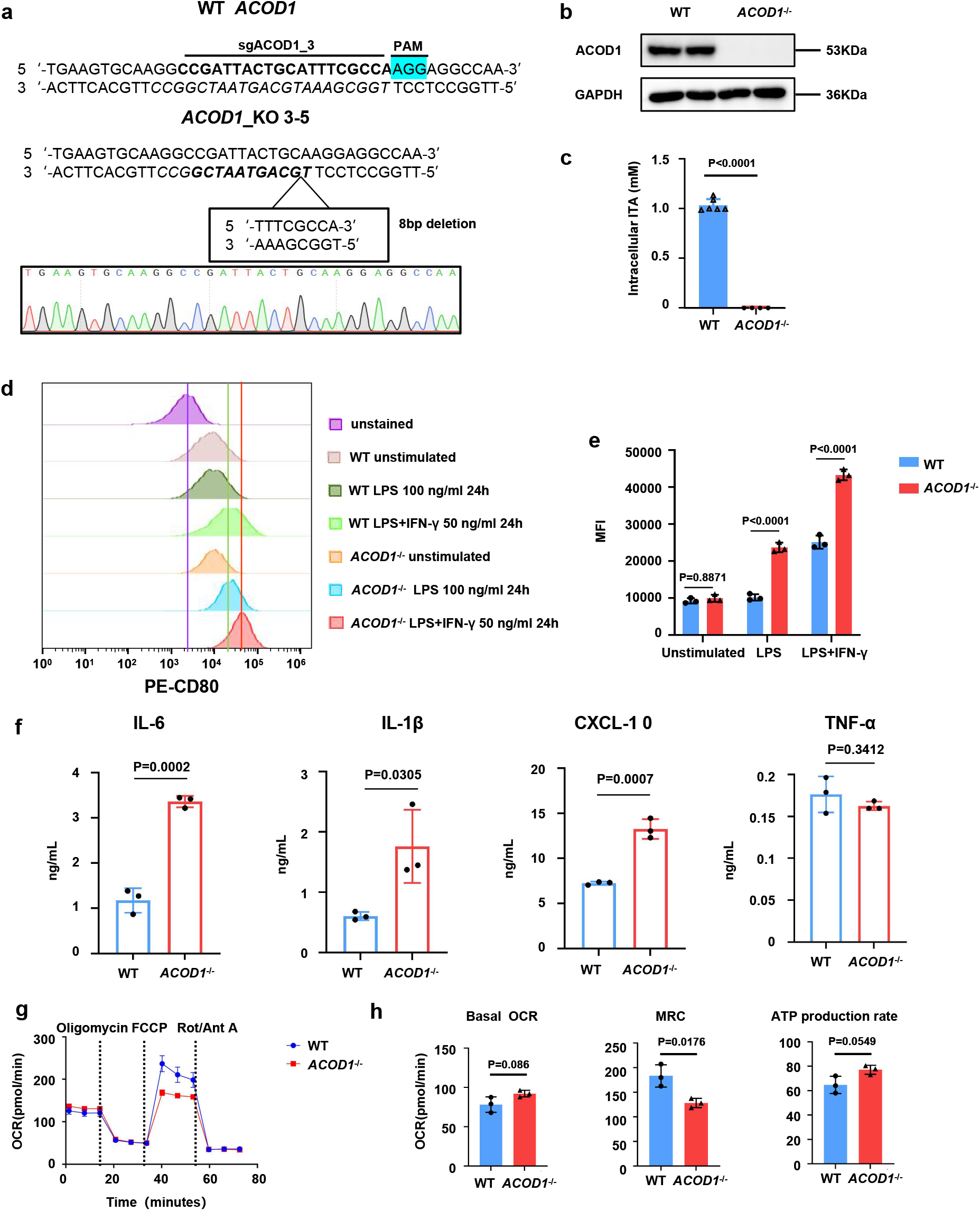
*ACOD1*-deleted human iPSC-derived macrophages demonstrated enhanced pro-inflammatory activation. **a,** Comparison of the DNA sequence in the *ACOD1* knockout iPSC clone (by Sanger sequencing) with the *ACOD1* WT DNA sequence showed an 8 bp deletion in the sgRNA targeted region. **b**, Western blotting for ACOD1 expression in WT and *ACOD1*^-/-^ iMACs after LPS and IFN-γ stimulation for 24 h. **c**, Mass spectrometry quantification of the cellular itaconate (ITA) concentration in WT and *ACOD1*^-/-^ iMACs after LPS and IFN-γ stimulation for 24 h (WT, n=6; *ACOD1*^-/-^, n=4). **d,e**, CD80 expression on WT and *ACOD1*^-/-^ iMACs and mean fluorescence intensity (MFI) quantification was determined by flow cytometry under different treatments, including 100 or 50 ng/mL LPS plus 50 ng/mL IFN-γ stimulation for 24 h (e, n=3). **f**, The levels of the indicated cytokines/chemokines in the medium of iMAC culturing were determined 24 h post IFN-γ and LPS challenge (n=3). **g**, Seahorse extracellular metabolic flux analysis of oxygen consumption rates (OCRs). LPS and IFN-γ stimulated WT or *ACOD1*^-/-^ iMACs were sequentially treated with oligomycin (1.5 µM), fluorcarbonylcyanide phenylhydrazone (FCCP; 2 µM), and rotenone and antimycin A (0.5 µM each) (n=3). **h**, Basal OCR, maximal respiration capacity (MRC), ATP production rate and spare respiration capacity (SRC) were calculated with Wave 2.4.0. (n=3 biological replicates representative of three independent experiments). **c, e, f, g and h**, Data was shown as mean ± SD. Statistics by two-way ANOVA test (e) or unpaired t test (c,f,h).

We then tried to examine whether *KEAP1* had the same effects on human iPSC-derived macrophages. We obtained an iPSC cell line with a 22 bp deletion on the *KEAP1* gene using the CRISPR-Cas9 technology (Extended Data Fig. 4a). KEAP1 protein expression was significantly decreased in the knockout cell line (Extended Data Fig. 4b). However, *KEAP1* was continuously expressed during the differentiation process from WT iPSCs to macrophages (Extended Data Fig. 4c), and we could not obtain differentiated macrophages from the *KEAP1* knockout iPSC line, suggesting *KEAP1* may be an essential gene in the process of macrophage differentiation.

**Fig. 4.**
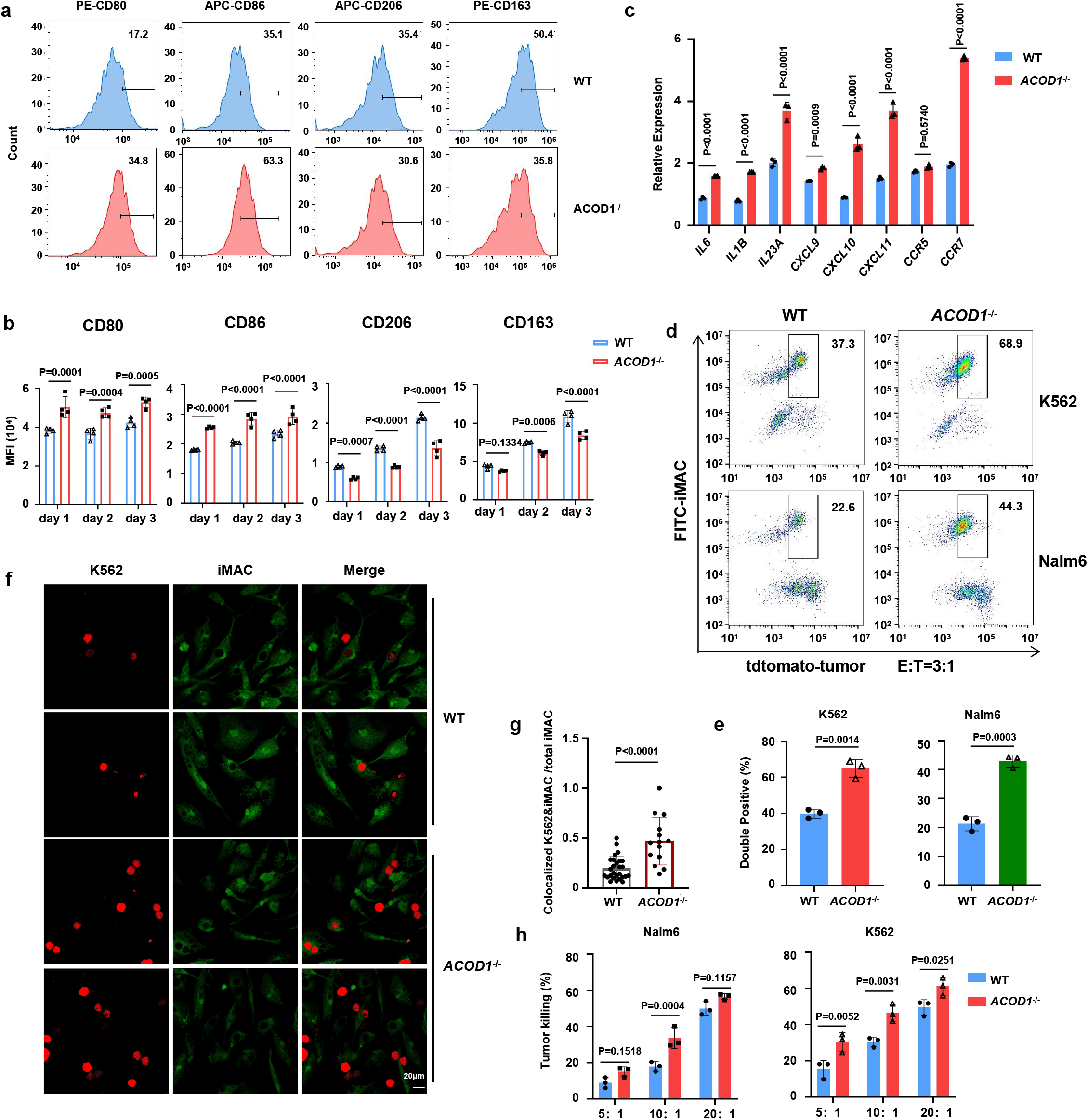
*ACOD1*^-/-^ iMACs had stronger phagocytosis and anti-cancer cell function. **a,** CD80, CD86, CD163, and CD206 expression in WT or *ACOD1*^-/-^ iMACs after co-cultured with Nalm6 (E: T=5:1) for 24 h were measured by flow cytometry and displayed as histograms. **b**, Quantification of MFI measured by flow cytometry after co-cultured with Nalm6 (E: T=5:1) for 24 h (day 1), 48 h (day 2), or 72 h (day 3) (n=4). **c**, qRT-PCR for mRNA expression of pro-inflammatory genes in WT and *ACOD1*^-/-^ iMACs after co-cultured with Nalm6 (E: T=5:1) for 24 h (n=3). **d,e**, Representative flow cytometry plots and quantification of double positive iMACs after WT and *ACOD1*^-/-^ iMACs were co-cultured with Nalm6 and K562 cells (E: T=3:1) for 24 h (e, n=3). **f,g**, Representative confocal images and quantification of K562 cells phagocytosed by WT or *ACOD1*^-/-^ iMACs after co-cultured for 24 h (g, WT, n=27; *ACOD1*^-/-^, n=14). The number of colocalized K562&iMAC and total iMAC in one view was used to calculate the ratio. **h**, Luciferase assays showing iMAC cytotoxicity against cancer cells when co-cultured with Nalm6 or K562 cells for 24 h (E: T=5:1, 10:1, or 20:1) (n=3). The luciferase gene has been introduced by lentivirus to tumor cells and expressed in tumor cells, so that tumor cell viability can be measured by D-luciferin sodium salt in a luciferase assay. **b, c, g, e and h**, Data was shown as mean ± SD. Statistics by two-way ANOVA test (b and h), unpaired t test (g and e).

### *ACOD1* deletion promoted pro-inflammatory activation in tMACs

The challenge of obtaining *KEAP1*-deleted iMACs enabled us to examine other players in the pathway. The KEAP1 protein can be modified and regulated via the alkylation of cysteine by a metabolite called itaconate^14^. Aconitate decarboxylase 1 (encoded by *ACOD1*) or Immune Responsive Gene 1 (IRG1) is the sole enzyme responsible for itaconate production and functions as an upstream regulator of KEAP1^15^. The alkylation of KEAP1 allows newly synthesized NRF2 to accumulate, transfer to the nucleus, and activate the transcription of anti-oxidant genes^16, 17^. According to this mechanism, we speculate *ACOD1* deletion may enhance the pro-inflammatory activation of macrophages, opposite to what KEAP1 does. Our CRISPR screen in iPSC-derived macrophages also identified sgRNAs targeting *ACOD1* enriched in the CD80-high population (Extended Data Fig. 5a). To further investigate the role of ACOD1 in human macrophage pro-inflammatory activation, we designed 4 sgRNAs targeting *ACOD1* (Extended Data Fig. 5b), T7 endonuclease assays revealed that sgRNA-2 and sgRNA-3 had higher cleavage activity (Extended Data Fig. 5c,d and e). We then generated *ACOD1*-deleted THP-1 cells in which the mRNA expression was significantly lower (Fig. 2a), and the protein expression of ACOD1 was nearly blank in sg*ACOD1*-2 and sg*ACOD1*-3 THP-1 derived macrophages (Fig. 2b). To reveal the function of ACOD1 in pro-inflammatory activation of macrophages, we found CD80 expression was higher in *ACOD1-*deleted macrophages after stimulation and the magnitude of difference was maintained after two days (Fig. 2c,d). The mRNA expression of pro-inflammatory genes showed an approximately 2-fold increase in *ACOD1-*deleted macrophages, such as *IL6* and chemokine genes *CXCL9*, *CXCL10,* and *CXCL11*, especially after 8 h of stimulation (Fig. 2e). When stimulated by LPS alone, about a 5-fold increase of CD80 expression could also be detected (Extended Data Fig. 5f,g). Collectively, these data demonstrate that *ACOD1* deletion promotes more sustainable pro-inflammatory activation of tMACs following stimulation.

**Fig. 5.**
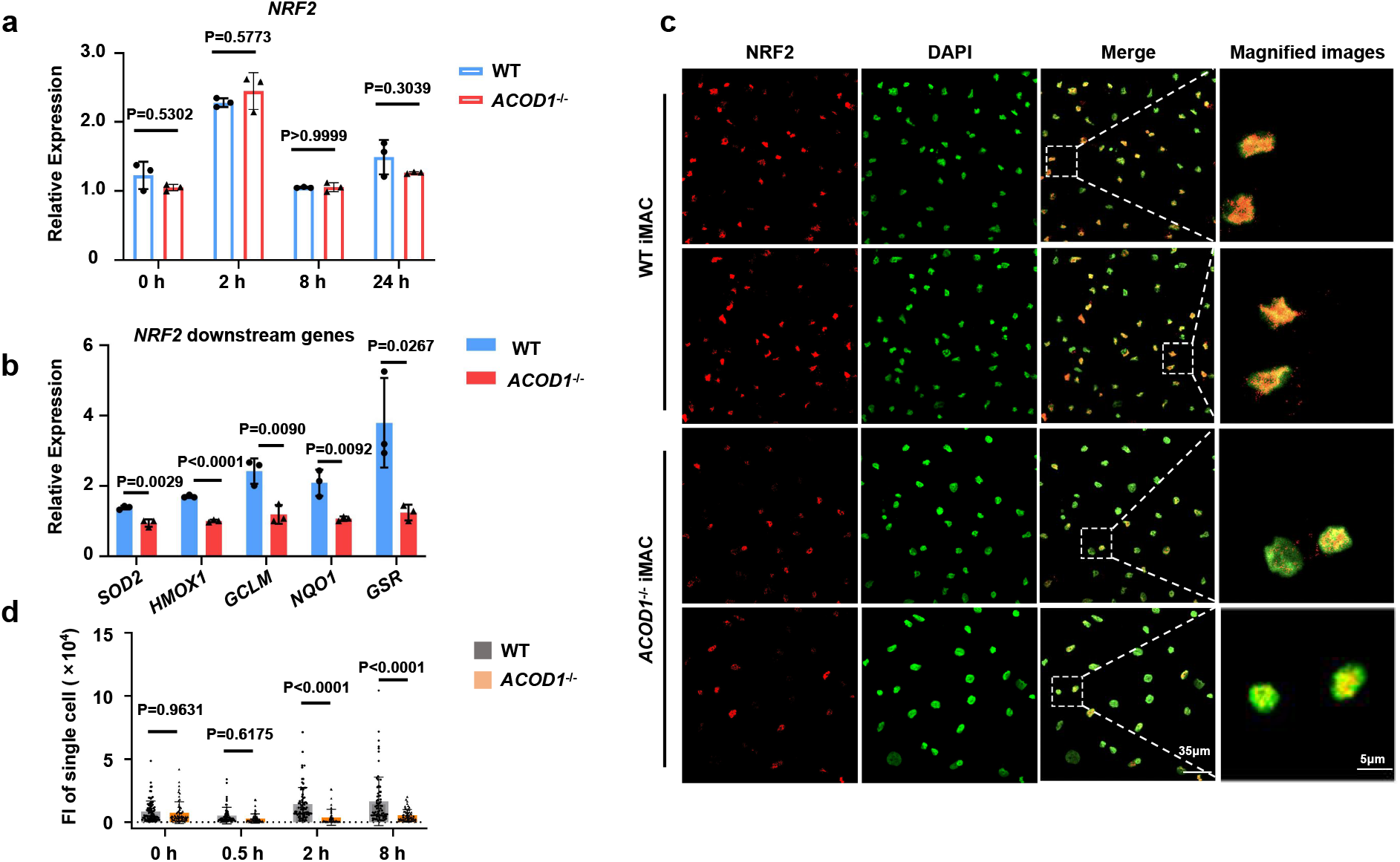
*ACOD1* deletion decreased nucleolar NRF2 protein expression and its activity in iMACs. **a**, qRT-PCR for mRNA expression of *NRF2* in WT and *ACOD1*^-/-^ iMACs after LPS and IFN-γ stimulation for 2, 8, or 24 h (n=3). **b**, qRT-PCR for mRNA expression of *NRF2* downstream genes in WT and *ACOD1*^-/-^ iMACs after LPS and IFN-γ stimulation for 24 h (n=3). **c**,**d**, Representative confocal images and quantification of the NRF2 protein in WT and *ACOD1*^-/-^ iMACs after LPS and IFN-γ stimulation for 2 h (d, n=60). **a,b** and **d,** Data was shown as mean ± SD. Statistics by two-way ANOVA test.

### *ACOD1-*deleted human iMACs demonstrated enhanced pro-inflammatory activation

To investigate whether *ACOD1* deletion contributes to pro-inflammatory activation in human iMACs, we knocked out *ACOD1* in human iPSC using the CRISPR/Cas9 technology with sgRNA-3. A new cell line with an 8 bp deletion on the fourth exon of the *ACOD1* gene was established (Fig. 3a). Differentiation from this engineered iPSCs to macrophages was successful, and the purity of macrophages reached to 96% on day 29 (Extended Data Fig. 6a,b). The deficiency of ACOD1 in mRNA (Extended Data Fig. 6c) and protein expression (Fig. 3b) was confirmed. As expected, the intracellular concentration of itaconate (ITA) was also significantly lower in *ACOD1* deficient iMACs (Fig. 3c). After 24 h of stimulation with LPS or LPS plus IFN-γ, the expression of CD80 was significantly higher in *ACOD1*^-/-^ iMACs (Fig. 3d,e). We further measured the mRNA expression of other pro-inflammatory genes to confirm this result. In line with elevated CD80 expression, pro-inflammatory genes *IL6, IL1B, IL1A, IL23A* and *CXCL-10* were also significantly higher in *ACOD1*^-/-^ iMACs (Extended Data Fig. 6d). We also validated the changes at the protein level with ELISA. Compared with WT iMACs, *ACOD1*^-/-^ iMACs had increased levels of pro-inflammatory cytokines and chemokines such as IL-6, IL-1β and CXCL-10 in the supernatant upon LPS and IFN-γ stimulation (Fig. 3f). To extend the finding that *ACOD1* restricted the iMAC pro-inflammatory state and the associated metabolic program, we measured real-time changes in cellular oxygen consumption (OCR) in WT and *ACOD1*^-/-^ iMACs. *ACOD1* deletion led to a decreased oxygen consumption rate (OCR) (Fig. 3g), including decreased maximal respiration capacity (MRC) (Fig. 3h), suggesting a decrease in mitochondrial function typically associated with the M2-like state in the absence of *ACOD1*. Together, these results demonstrate that *ACOD1* deletion promotes pro-inflammatory activation of iMACs, and decreases mitochondrial function upon pro-inflammatory stimulation.

**Fig. 6.**
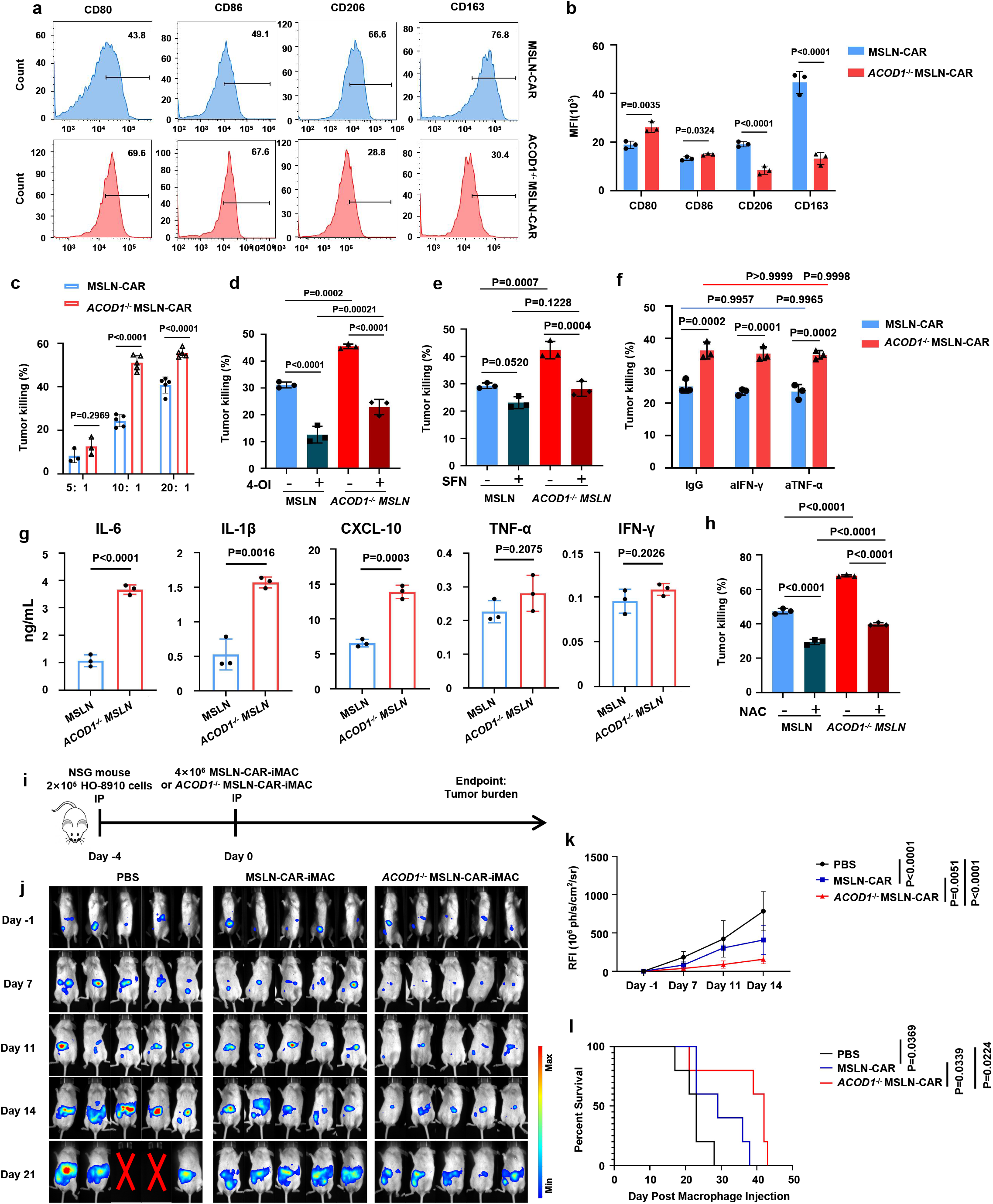
*ACOD1* deletion promoted anti-cancer cell activity of iMACs against solid tumors *in vitro* and *in vivo*. **a,b,** The expression and quantification of CD80, CD86, CD163, and CD206 in MSLN or *ACOD1*^-/-^ MSLN CAR-iMACs after co-cultured with HO-8910 cells (E: T=5:1) for 24 h were measured by flow cytometry and displayed as histograms (b, n=3). **c**, Luciferase assays for CAR-iMAC cytotoxicity activity against cancer cells when co-cultured with HO-8910 cells for 24 h (E: T=5:1, 10:1, or 20:1) (5:1, n=3; 10:1, n=5; 20:1, n=5). **d**, Luciferase assays for CAR-iMAC cytotoxicity activity against cancer cells with or without 4-OI addition when co-cultured with HO-8910 cells for 24 h (n=3) (E: T=10:1). iMACs were pre-treated with 4-OI (250 μM) or DMSO control for 3 h before challenge with LPS plus IFN-γ (50 ng/mL each) for 24 h. **e**, Luciferase assays for MSLN-CAR-iMAC cytotoxicity activity against cancer cells with or without SFN (10 μM) when co-cultured with HO-8910 cells for 24 h (E: T=10:1) (n=3). **f**, Luciferase assays for the cytotoxicity activity of the co-culture supernatant with IgG control, neutralizing antibody (10 μg/mL) of IFN-γ or TNF-α (n=3). The supernatant was collected after iMACs were co-cultured with HO-8910 cells for 24 h (E: T=10:1). **g**, The levels of the indicated cytokines/chemokines in the medium of iMAC-HO-8910 co-culture system were determined 24 h post IFN-γ and LPS challenge (n=3). **h**, Luciferase assays for MSLN-CAR-iMAC cytotoxicity activity against cancer cells with or without NAC (2.5 mM) when co-cultured with HO-8910 cells for 48 h (E: T=10:1) (n=3). (**b-h**) Data was shown as mean ± SD. (**b, c, d, e, f, h**) Statistics by two-way ANOVA test. **g**, Statistics by unpaired t test. **i**, A diagram of the in vivo treatment scheme. **j**, IVIS images showing progression of tumor in the above conditions (n=5 per group). **k**, Tumor burden on day -1, 7, 11, and 14 was quantified and displayed as mean ± SD. statistics by two-way ANOVA test. **l**, The Kaplan-Meier curve demonstrating survival of the mice. Statistics by two-tailed log-rank test.

### *ACOD1*^-/-^ iMACs demonstrated a stronger phagocytosis and anti-cancer cell function

To further investigate the role of ACOD1 in iMACs in the presence of tumor cells, Nalm6 or K562 cells were used to co-culture with WT iMACs or *ACOD1*^-/-^ iMACs. We found that, after co-culturing with Nalm6 cells for 24 h at an effector: target ratio of 5:1 or 3:1, the expression levels of M1-like markers CD80 and CD86 were higher in *ACOD1*^-/-^ iMACs, whereas M2-like markers CD163 and CD206 were lower (Fig. 4a and Extended Data Fig. 7a). Co-culturing with K562 cells had the similar results (Extended Data Fig. 7b,c). Importantly, a long-term co-culture assay revealed that the expression of M1-like markers remained elevated, whereas M2-like markers remained lower in *ACOD1*^-/-^ iMACs in three days (Fig. 4b), indicating that *ACOD1* deletion could contribute to a long-term maintenance of higher pro-inflammatory activation and resistance to conversion toward the anti-inflammatory state in the presence of Nalm6 tumor cells. In addition, mRNA expression of other M1-like marker genes was also significantly higher in *ACOD1*^-/-^ iMACs co-cultured with Nalm6 cells (Fig. 4c and Extended Data Fig. 7d), and their expression was also maintained higher over long term co-culturing (Extended Data Fig. 7e). Next, flow cytometry results support the stronger phagocytosis function of *ACOD1*^-/-^ iMACs against tumor cells (Fig. 4d,e). The isotype control and gating strategy of the phagocytosis assay was shown in Extended Data Fig. 7f-h. Confocal imaging analysis also showed that *ACOD1*^-/-^ iMACs co-cultured with K562 cells for 24 h had a stronger phagocytosis function (Fig. 4f,g). Finally, the luciferase assay showed *ACOD*1^-/-^ iMACs had higher cytolytic activity against tumor cells (Fig. 4h). Taken together, the above data demonstrate ACOD1 deletion promotes a stronger anti-tumor function upon tumor cell stimulation.

**Fig. 7.**
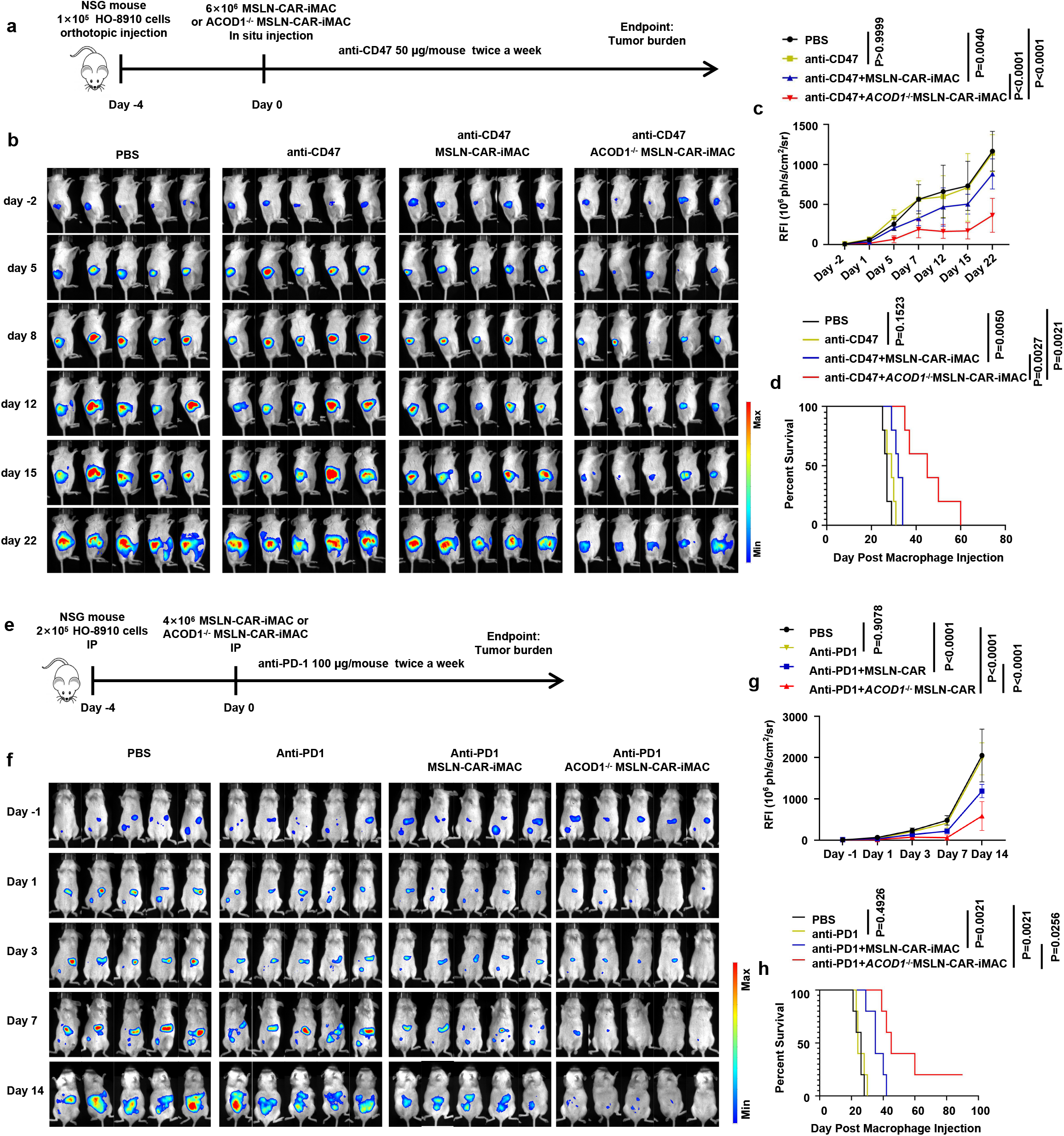
*ACOD1* deletion promoted the anti-ovarian cancer activity of iMACs combining with ICIs *in vivo*. **a,** A schematic of the in vivo study using HO-8910 cells for a mouse ovarian orthotopic injection model treated with MSLN-CAR-iMACs and *ACOD1*^-/-^ MSLN-CAR-iMACs, and combined with an anti-CD47 antibody. **b**, Tumor burden was determined by BLI. Images of representative time points were shown (n=5 per group). **c**, Quantification of tumor burden of representative time points was displayed as mean ± SD. Statistics by two-way ANOVA test. **d**, The Kaplan-Meier curve demonstrating survival of the mice. Statistics by two-tailed log-rank test. **e**, A schematic of the *in vivo* study using HO-8910 cells for a mouse intraperitoneal injection model treated with MSLN-CAR-iMACs and *ACOD1*^-/-^ MSLN-CAR-iMACs, and combined with an anti-PD1 antibody. **f**, Tumor burden was determined by BLI. Images of representative time points were shown (n=5 per group). **g**, Quantification of tumor burden of representative time points was displayed as mean ± SD. Statistics by two-way ANOVA test. **h**, The Kaplan-Meier curve demonstrating survival of the mice. Statistics by two-tailed log-rank test.

### *ACOD1* deletion decreased the expression of the nuclear NRF2 protein and its activity in iMACs

It was demonstrated that itaconate was a crucial anti-inflammatory metabolite that acts via NRF2^14^. To understand the molecular mechanisms of ACOD1 depletion in our iMAC system, we examined NRF2 and its downstream genes in iMACs. We found mRNA expression of *NRF2* had no significant difference in WT and *ACOD1*^-/-^ iMACs (Fig. 5a). However, the expression of *NRF2* downstream genes decreased significantly in *ACOD1*^-/-^ iMACs, such as *SOD2, HMOX1, GCLM, NQO1,* and *GSR* (Fig. 5b). Confocal imaging showed that the total NRF2 protein level in the nucleus decreased significantly in *ACOD1*^-/-^ iMACs, especially after LPS and IFN-γ stimulation for 2 and 8 h (Fig. 5c, d and Extended Data Fig. 8a-c). One of the NRF2 targets is TNFAIP3 (A20) which is a negative regulator of the NF-κB pathway and macrophage activation^18^. We measured *A20* expression in iMACs and found that it decreased in *ACOD1*^-/-^ iMACs (Extended Data Fig. 8d), which is likely to mediate increased NF-κB activity. To further validate the functional effect of NRF2 on macrophage activation, we designed three gRNAs targeting *NRF2* (Extended Data Fig. 9a), which all successfully lowered the mRNA expression of *NRF2* (Extended Data Fig. 9b). Depletion of NRF2 recapitulated *ACOD1* deletion in that CD80 expression was higher in sg*NRF2*s transduced cells compared to WT controls after LPS and IFN-γ stimulation for 24 h (Extended Data Fig. 9c), and consistently mRNA expression of pro-inflammatory genes was also higher (Extended Data Fig. 9d). Together, these results demonstrate that *ACOD1* deletion decreased NRF2 activity to allow pro-inflammatory activation of iMACs.

### *ACOD1* deletion promoted anti-cancer cell activity against solid tumors of CAR-iMACs *in vitro* and *in vivo*

Adoptive cell therapy with genetically modified immune cells has been established as a promising approach for cancer treatment. However, applications to solid tumors have proven challenging. To improve the anti-solid tumor functions of iMACs, we used our previously established CAR-iMAC system, in which we stably expressed the first generation of anti-mesothelin (MSLN) CAR with CD3ζ as the intracellular domain in human iPSCs and differentiated them to produce MSLN-CAR-iMACs to kill mesothelin-expressing ovarian tumors both *in vitro* and *in vivo*^10^. We then performed a detailed comparison of MSLN-CAR-iMACs and *ACOD1*^-/-^ MSLN-CAR-iMACs. *ACOD1*^-/-^ MSLN-CAR-iMACs expressed more pro-inflammatory marker proteins (CD80 and CD86) after being co-cultured with HO-8910 ovarian cancer cells for 24 h (Fig. 6a,b). Lower anti-inflammatory marker proteins (CD163 and CD206) were consistently detected in *ACOD1*^-/-^ MSLN-CAR-iMACs (Fig. 6a,b). *In vitro* tumor cell killing assay revealed that *ACOD1*^-/-^ MSLN-CAR-iMACs significantly increased anti-tumor activity (Fig. 6c), which could be dampened by supplementing a cell permeable 4-Octyl Itaconate (4-OI)^19^ (Fig. 6d). To further dissect the downstream signaling related to NRF2, a chemical NRF2 activator sulforaphane (SFN)^20^ was added to the co-culture system, which abrogated the enhanced capacity in *ACOD1*^-/-^ MSLN-CAR-iMACs (Fig. 6e). To examine the cytolytic mechanisms in addition to phagocytosis, we collected the supernatant from the co-culture of tumor cells and CAR-iMACs, and then added it to another well of tumor cells. To our surprise, the supernatant alone had tumor killing function (Fig. 6f), which was not abrogated by neutralizing antibodies of TNF-α and IFN-γ (Fig. 6f), two cytolytic cytokines. Interestingly, compared with MSLN-CAR-iMACs, *ACOD1*^-/-^ MSLN-CAR-iMACs had increased inflammatory cytokines such as IL-6, IL-1β and CXCL-10, but not TNF-α and IFN-γ when co-cultured with HO-8910 cells for 24 h (Fig. 6g). Then we explored other potential cytolytic factors in the medium. It has been reported that ROS is produced upon LPS stimulation and through TLR^21^, and it can be dampened by supplementing itaconate in macrophages^14^. Compared to MSLN-CAR-iMACs, ROS production was also elevated in *ACOD1*^-/-^ MSLN-CAR-iMACs (Extended Data Fig. 10a, b). To examine the function of ROS in tumor killing ability of *ACOD1*^-/-^ MSLN-CAR-iMACs, we added the anti-oxidant reagent N-Acetyl-L-cysteine (NAC) to eliminate ROS in the tumor-iMAC co-culture system. The tumor killing capacity of CAR-iMACs was significantly blocked by NAC (Fig. 6h). This result demonstrated that ROS contributed to the enhanced tumor killing ability of *ACOD1*^-/-^ MSLN-CAR-iMACs.

To evaluate the anti-tumor activity of *ACOD1*^-/-^ MSLN-CAR-iMACs *in vivo*, we used two different xenograft solid tumor models with the NSG mice. In the first model, the mice were inoculated intraperitoneally (IP) with luciferase-expressing HO-8910 cells. After four days, the mice received a single IP injection of iMACs (Fig. 6i), and were monitored by bioluminescent imaging (BLI) afterwards (Fig. 6j). Compared with the untreated or the MSLN-CAR-iMACs treated mice, treatment with *ACOD1*^-/-^ MSLN-CAR-iMACs led to significant inhibition of tumor growth (Fig. 6k). This improved anti-tumor activity also led to markedly improved survival time (Fig. 6l). Then we examined the pro-inflammatory activity of iMACs *in vivo*. iMACs were injected intratumorally, and the pro-inflammatory markers were significantly elevated in *ACOD1*^-/-^ MSLN-CAR-iMACs compared with unmodified MSLN-CAR-iMACs after injection for 7 days (Extended Data Fig. 11a, b) or 14 days (Extended Data Fig. 11c, d). These results indicated that *ACOD1-*depleted CAR-iMACs could keep an enhanced pro-inflammatory activity *in vivo* for at least 14 days.

Consistent results were obtained in another setting of pancreatic cancer. The M1 markers were significantly elevated in *ACOD1*^-/-^ MSLN-CAR-iMACs compared with unmodified MSLN-CAR-iMACs after co-cultured with AsPC-1 pancreatic cancer cells for 24 h (Extended Data Fig. 12a). *In vitro* tumor killing capacity was enhanced in *ACOD1*^-/-^ MSLN-CAR-iMACs (Extended Data Fig. 12b) which could be reversed by supplementing 4-OI (Extended Data Fig. 12c). The expression of pro-inflammatory genes was also elevated in *ACOD1*^-/-^ MSLN-CAR-iMACs (Extended Data Fig. 12d). In line with the *in vitro* results, *in vivo* assay using a pancreatic tumor mouse model with intraperitoneally injected AsPC-1 cells also demonstrated the stronger anti-tumor activity of *ACOD1*^-/-^ MSLN-CAR-iMACs (Extended Data Fig. 12e-h).

### *ACOD1* deletion promoted the anti-tumor activity of MSLN-CAR-iMACs in combination with immune check point inhibitors *in vivo*

Tumor cells evade normal immune system via transmitting inhibitory signals to myeloid cells^22^ and lymphocytes^23^. Immune checkpoint is one of the mechanisms that regulate cancer immune escape. For instance, CD47 is expressed on many cancer cells, and binding of CD47 to signal-regulatory protein α (SIRPα) on macrophages results in inhibition of macrophage phagocytic activity^24^. Programmed cell death protein 1 (PD-1) is an immune checkpoint receptor mainly upregulated on activated T cells for the induction of immune tolerance. It’s well known that PD-1-PD-L1 blockade could activate T cells^25^. PD-1 is also expressed on tumor associated macrophages, and its expression is negatively correlated with phagocytic potency of macrophages^26^. We hypothesized that the combination of CAR-iMACs with ICIs may enhance the anti-tumor activity. So we assessed two combination immunotherapy strategies using MSLN-CAR-iMACs with the anti-CD47 antibody and the anti-PD-1 antibody, respectively.

In the first xenograft model, HO-8910 cells were inoculated through orthotopic injection at the ovary of the mice. After four days, the mice received a single in situ intratumoral injection of iMACs. At the same time, the mice received IP injections of a low dose anti-CD47 antibody, and it was kept twice a week to further enhance the function of CAR-iMACs by blocking the “don’t eat me” signal (Fig. 7a). The tumor growth was monitored by BLI (Fig. 7b). Compared with untreated tumor-bearing mice, the low-dose anti-CD47 antibody treatment alone could not inhibit tumor growth. The combination of the low-dose anti-CD47 antibody with the MSLN-CAR-iMACs could inhibit tumor growth to some extent. Importantly, the combination of low-dose anti-CD47 antibody with *ACOD1*^-/-^ MSLN-iMACs had the most superior tumor suppression effect (Fig. 7c). This improved anti-tumor activity led to markedly improved survival time compared with all other conditions (Fig. 7d). In the second xenograft model with the anti-PD-1 antibody, the mice were inoculated intraperitoneally (IP) with luciferase-expressing HO-8910 cells. After four days, the mice received a single IP injection of iMACs. Meanwhile, the mice received IP injections of a low dose of the anti-PD-1 antibody, and subsequently the antibody was used twice a week to further block the PD-1-PD-L1 axis (Fig. 7e). The mice were monitored by bioluminescent imaging afterwards (Fig. 7f). Compared with other groups, the combination of the low-dose anti-PD1 antibody with *ACOD1*^-/-^ MSLN-CAR-iMACs had the most superior tumor suppression effect (Fig. 7g), and markedly lengthened the survival time (Fig. 7h). Together, these data strongly demonstrated that *ACOD1*-deleted CAR-iMACs combined with ICIs had the most superior anti-tumor activity.

## Discussion

The ACOD1/KEAP1/NRF2 axis plays a crucial role in maintaining redox balance and macrophage polarization in mouse and human macrophages^14^. In the mouse sepsis syndrome model, *Keap1* deletion in macrophages resulted in reduced levels of inflammatory mediators, organ injury, bacteremia and mortality, whereas Nrf2 deletion had the opposite effects^27^. We found that KEAP1 plays an important role in macrophage pro-inflammatory activity through a pooled CRISPR screen of metabolic genes. Our study elucidated that *KEAP1* deletion inhibited whereas ACOD1 deletion promoted macrophage pro-inflammatory activity through regulating NRF2. We used screens in both THP1 and iMACs, and the limitation for the screen in iMACs was that many metabolic genes are necessary for macrophage differentiation and survival, and thus the essential genes will be missed in the list of positive candidates. In our case, only several genes were identified from the iMAC screen, including *ACOD1*. The reason that ACOD1 can be picked up might be that its expression is not required in unstimulated macrophages or during macrophage differentiation, and it is only induced upon LPS+IFN-γ stimulation. Thus it is not considered as an essential gene. To obtain more candidates, an inducible system of CRISPR screen would be a better choice in which Cas9 can be induced after differentiation^28–30^. Besides *ACOD1*, many other highly ranked genes coming out of our metabolic screen may also contribute to macrophage pro-inflammatory activity, such as *ULK1*, *GCLM*, *PPARD*, *GPD2,* and so on. GPD2 regulates LPS-induced macrophage tolerance via a pathway distinct from ACOD1^31^. Therefore, the double knockout of two genes that work in orthogonal pathways may represent new metabolic engineering strategies to further enhance macrophage functions in cancer immune cell therapies.

ACOD1 plays a crucial role in mitochondrial metabolism, which is tightly connected to many aspects of cellular functions. We also observed the maximal oxygen consumption was decreased in *ACOD1*^-/-^ iMACs. ACOD1 produces itaconate in response to pathogen infection and inflammation^15^. Itaconate can inhibit inflammasome activation^32, 33^ or regulate immune tolerance through succinate dehydrogenase (SDH) in macrophages^34–36^. For instance, Lampropoulou et.al showed itaconate could inhibit SDH and resulted in increased succinate level, and Irg1 deletion led to abrogation of succinate accumulation^36^, and Chen et.al showed overexpression of Irg1, but not its catalytically inactive mutant, results in elevated intracellular levels of ITA and succinate^37^. Thus we can not exclude that ACOD1 deletion might influence the macrophage polarization phenotype through other downstream metabolites.

Since ACOD1 and itaconate have important roles in the anti-inflammatory effects of macrophages, most previous studies focused on their functions in infectious diseases^37, 38^, However, limited studies have shown the role of human ACOD1 in immune-oncology and its possible applications in myeloid cell-based adoptive cell transfer in cancer immunotherapy. Engineered iMACs such as CAR-iMACs provide a new platform for cancer immune cell therapy^9, 39^. In this study, we demonstrated ACOD1 deficiency could promote stronger anti-solid tumor function than wild type MSLN-CAR-iMACs. We have also shown that *ACOD1*^-/-^ MSLN-CAR-iMACs combined with low dose ICIs could further elevated anti-ovarian cancer capacity. Thus, ACOD1 is a new metabolic target to engineer CAR-iMACs, in order to elevate their anti-tumor function and to eliminate tumor cells.

Macrophages can both kill cancer cells and modulate the tumor microenvironment depending on their phagocytosis function and pro-inflammatory activity^40^, which can be greatly enhanced by ACOD1 deletion as we have shown above in this work. Our results also showed that the iMAC-tumor co-culture supernatant alone indeed had tumor killing function as well. Either cytokines with cytolytic activity or ROS in the supernatant could confer the function. Neutralizing antibodies of TNF-α and IFN-γ did not abrogate tumor killing function of the supernatant (Fig. 6f). However, we could not completely rule out there could be other cytokines that mediated the cytolytic activity. Supplementing NAC reversed the phenotype, suggesting ROS contributed to the tumor killing function. Regarding indirect functions through other immune cells, as our experiments were conducted either in the absence of other immune cell types *in vitro*, or based on immuno-deficient NSG mice, which do not have the endogenous NK and T cells, the tumor killing activity in these settings was unlikely through stimulating NK and T cells. We can not exclude that in a humanized model, the engineered CAR-iMAC cells may also influence the other endogenous immune cells to confer their anti-tumor activity. Overall, our current data support a multi-level mechanism of CAR-iMACs in tumor killing activity, including enhanced direct phagocytosis and more ROS produced by *ACOD1* knockout.

Cytokines secreted from highly proliferative immune cells might lead to cytokine release syndrome. However, as macrophages do not have the capacity to proliferate *in vivo*, the amount of secreted cytokines might not reach to a level that can lead to toxicity in patients. Nevertheless, the effect of increased pro-inflammatory cytokines in *ACOD1*^-/-^ macrophages *in vivo* merits further investigations using humanized models to provide guidance for choosing the optimal dose in clinical research in the future.

## Methods

### Cell Lines

THP-1 cells, HO-8910 cells, K562 cells, Nalm6 cells, and 293T cells were obtained from the National Collection of Authenticated Cell Cultures and cultured according to standard protocols. Human iPSCs were obtained from the reprogramming of peripheral blood mononuclear cells from a volunteer donor, as described before ^10^. Human iPSCs were cultured in mTeSR medium (85852, STEMCELL Technologies) with Matrigel Matrix (354277, Corning) coated plates.

### Plasmid construction and single guide RNA cloning

All the Cas9-expressing THP-1 cells or iPSC lines in this study were derived by lentiviral transduction with a Cas9 expression vector containing an optimized sgRNA backbone (LentiCRISPR v2; Addgene, 52961). All of the sgRNAs were cloned into the LentiCRISPR v2 vector following the protocol described before^41^. The annealed sgRNA oligonucleotides were ligated with T4 DNA ligase (M0569S, NEB) to the BsmB1-digested LentiCRISPR v2 vector.

### Lentivirus production

We produced lentivirus using HEK293T cells cultured in DMEM supplemented with 1% penicillin-streptomycin and 10% FBS. The CRISPR library vectors (Human CRISPR Metabolic Gene Knockout Library; Addgene, Pooled Library #110066) ^12^ or the single sgRNA vectors, envelop vector pMD2.G, and packaging vector psPAX2 were mixed in a 4:3:1 ratio in OPTI-MEM (Thermo Fisher Scientific, 31985070) and PEI (Polysciences, 9002-98-6), and transfected into HEK293T cells at 80% to 90% confluence in 10-cm tissue culture plates. The supernatant was collected at 24, 48, and 72 h post-transfection, filtered via a 0.45 µm filtration unit (Millipore, Cat# SLHVR33RB), and mixed overnight at 4 °C with one-third volume of 30% PEG8000. The medium was concentrated at 4200 rpm for 30 min at 4 °C. The pellet was resuspended in PBS and stored at −80 °C.

### Transduction of lentivirus containing sgRNAs

For transfection of THP-1 cells and iPSCs, we infected cells with lentivirus and 5 µg/mL polybrene overnight, and the medium was changed the following day. After puromycin (1 µg/mL for THP-1 cells and 250 ng/mL for iPSC) selection for seven days, >95% of the population was transfected, and the cells were ready to be used for the subsequent experiments.

### Pooled CRISPR screen

1.5×10^7^ THP-1 cells were transduced with a viral library for 24 h (MOI = 0.3). After puromycin (1 µg/mL) selection for seven days, 1.5×10^7^ transduced cells were collected as input samples. The other transduced cells were treated with PMA (50 ng/mL) for 48 h, then stimulated by LPS (50 ng/mL) plus IFN-γ (50 ng/mL) for 24 h. The stimulated cells were harvested and stained with anti-human CD80-FITC (305206, BioLegend) for 15 min at room temperature. The CD80-high and CD80-low cells were separated by flow cytometry sorting. The genomic DNA of cells was isolated, and the sgRNA library was barcoded and amplified for two rounds of PCR. PCR products were purified for sequencing on an Illumina HiSeq. The sequencing data was analyzed by MAGeCK^42^.

### Generation of CRISPR/Cas9 knockout cells

LentiCRISPR v2 vectors targeting *KEAP1* and *ACOD1* were constructed as described before^41^. The THP-1 cells and iPSC were infected with lentivirus expressing Cas9 and sgRNAs targeting *KEAP1* and *ACOD1*. After puromycin selection for seven days, the THP-1 cells were expanded, and knockout efficiency was verified using qPCR and western blotting. After puromycin selection for three days, iPSCs were passaged, and the clones grown from single cells were picked up and expanded. The knockout efficiency of iPSC was verified by sequencing, qPCR, and western blot analyses.

### Derivation of iMACs from iPSCs

The derivation of iMAC from iPSCs has been previously described^10^. Briefly, 8000 iPSCs were seeded in 96-well round-bottom plates with APEL2 medium (05271, STEMCELL Technologies) containing 100 ng/mL human Stem Cell Factor (SCF), 50 ng/mL human Vascular Endothelial Growth Factor (VEGF), 10 ng/mL recombinant human Bone Morphogenetic Protein 4 (BMP-4), 5 ng/mL human FGF-basic (154 a.a.), and 10 mM Rho kinase inhibitor (ROCK inhibitor, Y27632, Sigma). After eight days of hematopoietic differentiation, spin embryoid bodies (EBs) were transferred into Matrigel-coated 6-well plates under macrophage differentiation conditions. Macrophage differentiation medium is StemSpan-XF (100-0073, STEMCELL Technologies) containing 10 ng/mL human FGF-basic (154 a.a.), 50 ng/mL human Vascular Endothelial Growth Factor (VEGF), 50 ng/mL human Stem Cell Factor (SCF), 10 ng/mL recombinant human Insulin-like Growth Factor-1 (IGF1), 20 ng/mL IL-3, 50 ng/mL recombinant human M-CSF, and 50 ng/mL recombinant human GM-CSF. The floating cells were collected from the supernatant and directly transferred into uncoated 6-well plates in macrophage culture medium. The macrophage culture medium is StemSpan-XF containing 50 ng/mL recombinant human M-CSF and 50 ng/mL recombinant human GM-CSF.

### Flow cytometry

The tMACs or iMACs were stimulated with LPS and IFN-γ for the indicated time. The single-cell suspensions were then prepared and incubated with an antibody or antibody cocktails for 15 min at room temperature for cell surface staining. Antibodies used in this study were PE Human IgG1 Isotype Control (403503, Biolegend), APC Human IgG1 Isotype Control (403505, Biolegend), FITC Human IgG1 Isotype Control (403507, Biolegend), APC anti-human CD206 (321109, Biolegend), APC anti-human CD86 (374207, Biolegend), PE anti-human CD80 (305208, Biolegend), PE anti-human CD163 (333605, Biolegend), FITC anti-human CD14 (301803, Biolegend) and APC anti-humanCD11B (301309, Biolegend). Data were recorded on Beckman DxFLEX and analyzed with the FlowJo V10 software.

### Enzyme-linked immunosorbent assay

The supernatant of iMAC culture or tumor-iMAC co-culture was collected and centrifuged at 300×g for 10 minutes to remove the precipitate. Human IL-6, IL-1β, CXCL-10, IFN-γ and TNF-α were quantified using Elisa kits (MultiSciences, EK106, EK101B, EK168, EK180, EK182) following the manufacturer’s protocols.

### *In vivo* anti-tumor assay

For *in vivo* experiments, 6–10-week-old female NOD-scid IL2Rg_null_ (NSG) mice(Gempharmatech, Jiangsu, n=5 per group) were used. All mice were maintained under pathogen-free conditions under the Zhejiang University Institutional Animal Care and followed the committee’s approved protocols. In the first ovarian cancer mouse model, 2×10^5^ luciferase gene expressing HO-8910 cells were inoculated intraperitoneally (IP) before treatment (day -4). After tumor cell inoculation, mice were randomly assigned to experimental groups. Four days later, 4×10^6^ MSLN-CAR-iMACs or ACOD1^-/-^ MSLN-CAR-iMACs were inoculated IP (day 0) for therapy. The tumor burden was determined by bioluminescence imaging (BLI) using an IVIS Imaging System (Biospace Lab).

In the ovarian cancer orthotopic injection mouse model with CAR-iMAC and anti-CD47 antibody combined therapy, 1×10^5^ luciferase gene expressing HO-8910 cells were inoculated directly into ovary before treatment (day -4). After tumor cell inoculation, mice were randomly assigned to experimental groups. Four days later, mice received a single in situ injection of 6×10^6^ MSLN-CAR-iMAC or ACOD1^-/-^ MSLN-CAR-iMACs (day 0) combined with a low-dose CD47 antibody (50 µg/mouse, twice a week) for therapy. Tumor burden was determined by BLI.

In ovarian cancer mouse model with CAR-iMAC and the anti-PD1 antibody combined therapy, 2×10^5^ luciferase gene expressing HO-8910 cells were inoculated intraperitoneally (IP) before treatment (day -4). After tumor cell inoculation, mice were randomly assigned to experimental groups. Four days later, mice received a single injection of 4×10^6^ MSLN-CAR-iMAC or ACOD1^-/-^ MSLN-CAR-iMACs intraperitoneally (day 0) combined with a low-dose anti-PD1 antibody (100 µg/mouse, twice a week) for therapy. The tumor burden was determined by bioluminescence imaging (BLI) using an IVIS Imaging System.

In the pancreatic cancer mouse model, 6–10-week-old male NSG mice(n=5 per group) were used. 1×10^5^ AsPC-1 cells were inoculated intraperitoneally before treatment (day-4). After tumor cell inoculation, mice were randomly assigned to experimental groups. AsPC-1 cells grow fast in vivo, in order to get a better therapeutic effect, a higher effect target ratio was used. Four days later, 1.5 ×10^7^ MSLN-CAR-iMACs or ACOD1^-/-^ MSLN-CAR-iMACs were intraperitoneally injected (day 0). The tumor burden was determined by BLI later.

### Western blotting

Pellets from 1×10^6^ cells were collected and resuspended with 100 µL RIPA Buffer (Beyotime, Cat# P0013J). The samples were incubated on ice for 30 min and centrifuged at 13000 rpm for 15 min at 4 °C. The supernatant was collected, and the protein concentration was measured by BCA analysis (Thermo Scientific, Cat# 23225). Approximately 50 µg of total protein was loaded for western blotting.

### Real-time reverse transcription-PCR

RNA was extracted from macrophages or tumor cells using Total RNA Isolation Kit V2 (Vazyme, Cat# RC112-01). Reverse transcription from RNA to cDNA use Hiscript Reverse Transcriptase (Vazyme, Cat# R302-01). PCR reactions were performed on a CFX96 Real-Time PCR System (Bio-Rad Laboratories) using ChamQ Universal SYBR qPCR Master Mix (Vazyme, Cat# Q711-02).

### Metabolic studies

Oxygen consumption rate (OCR) was measured using a Seahorse XF Cell Mito Stress Test Kit (Agilent, 103015-100). iMACs were resuspended in an RPMI1640 medium containing LPS (50 ng/mL) plus IFN-γ (50 ng/mL) and then seeded at 5×10^4^ cells/well in an XF96 plate. Eight hours later, the RPMI1640 medium was changed to XF RPMI medium. The oxygen consumption rate was measured (pmol/min) in real-time in an XFe96 Extracellular Flux Analyzer. iMACs were stimulated with LPS and IFN-γ for 24 h, and the OCR was in response to 1.5 µM oligomycin, 2 µM fluorcarbonylcyanide phenylhydrazone (FCCP) and 500 nM rotenone and antimycin A. Basal OCR, maximal respiration capacity (MRC), ATP linked respiration, and mitochondrial spare respiratory capacity (SRC) was calculated by WAVE V2.6 software.

### RNA-seq

Total RNA was isolated and purified using FastPure Cell/Tissue Total RNA Isolation Kit V2 (Vazyme, RC112-01) from 2×10^6^ tMACs according to the manufacturer’s protocol. RNA qualification was performed using Nanodrop to check RNA purity (OD260/0D280) and Agilent 2100 to check RNA integrity. A total amount of 2 µg RNA per sample was used for RNA-seq libraries preparation. RNA-seq libraries were prepared using VAHTS Stranded mRNA-seq Library Prep Kit for Illumina V2 (Vazyme, NR612-02) according to the manufacturer’s protocol and sequenced on an Illumina Hiseq 2500. The threshold of differentially expressed genes is p-adj < 0.05. The color descending from red to blue in the heatmaps of differentially expressed genes indicated log10 (FPKM+1) from large to small.

### Gene set enrichment analysis

To identify “biological signatures” depleted or enriched following CD80-based sorting or in the *KEAP1* knockout macrophages, we used DAVID Bioinformatics Resources (https://david.ncifcrf.gov/). We focused on the biological oncology of the GO gene sets to obtain the indicated enrichment score.

### Statistical analysis

All data are presented as mean ± SD. Comparisons between different groups were analyzed by the one-way analysis of variance (ANOVA), two-way analysis of variance (ANOVA), and unpaired two-tailed Student’s *t*-test. Kaplan-Meier survival curves were compared with the log-rank test. Statistical analyses were performed in GraphPad Prism 9.0.0 software using the statistical tests indicated for each experiment. All tests were considered significant at p < 0.05.

### Data availability

The RNA-seq data that support the findings of this study have been deposited in the GEO under the following accession codes: GSE216352. The CRISPR Screen datasets have been deposited in the GEO under the accession number GSE216353. All other data supporting the findings of this study are available from the corresponding author on reasonable request. Source data are provided with this paper.

## Acknowledgments

We thank Tiffany Horng (School of life science and technology, ShanghaiTech University) for providing advises on this work. This work was sponsored by the National Key Research and Development Program of China (2018YFA0107103 and 2018YFC1005002), the National Natural Science Foundation of China (31871453 and 91857116), Zhejiang Innovation Team grant (2019R01004), and the Zhejiang Natural Science Foundation (LR19C120001).

---

**Extended Data Fig. 1.**
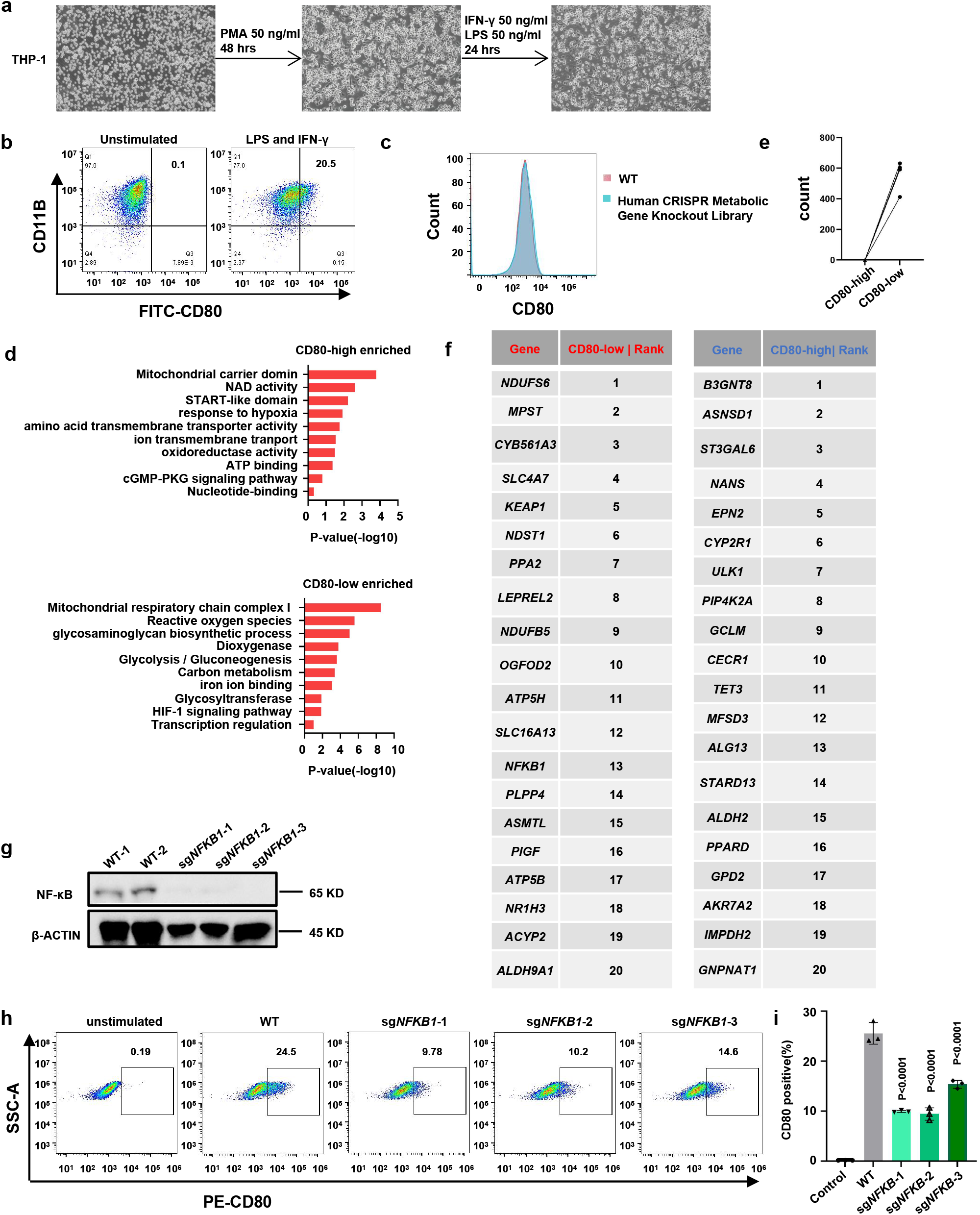
Identifying the metabolic genes involved in human macrophage activation, related to **Fig. 1**. **a**, Microscopic pictures showing THP-1 cell differentiation and polarization. **b**, Flow cytometry plots and percentage of CD80 expression on WT tMACs with or without LPS and IFN-γ stimulation for 24 h. **c**, WT and human Metabolic Gene CRISPR Library virus-infected THP-1 cells were differentiated into macrophages, and CD80 expression was measured by flow cytometry and demonstrated as histograms. **d**, GO term enrichment analysis with enriched sgRNA-targeted genes in CD80-high population (up), and CD80-low population (down). **e**, Counts of sgRNAs targeting *KEAP1* detected in the CD80-high and CD80-low samples. **f**, Top 20 sgRNA-targeted genes enriched in the CD80-low populations and CD80-high populations identified by the CRISPR Screen in tMACs. **g**, The protein level of NF-κB in WT and *NFKB1*-depleted THP-1 cells. **h,i**, Flow cytometry plots and quantification of CD80 expression on unstained, unstimulated, WT and *NFKB1*-depleted tMACs (i, n=3). Statistics by one-way ANOVA test.

**Extended Data Fig. 2.**
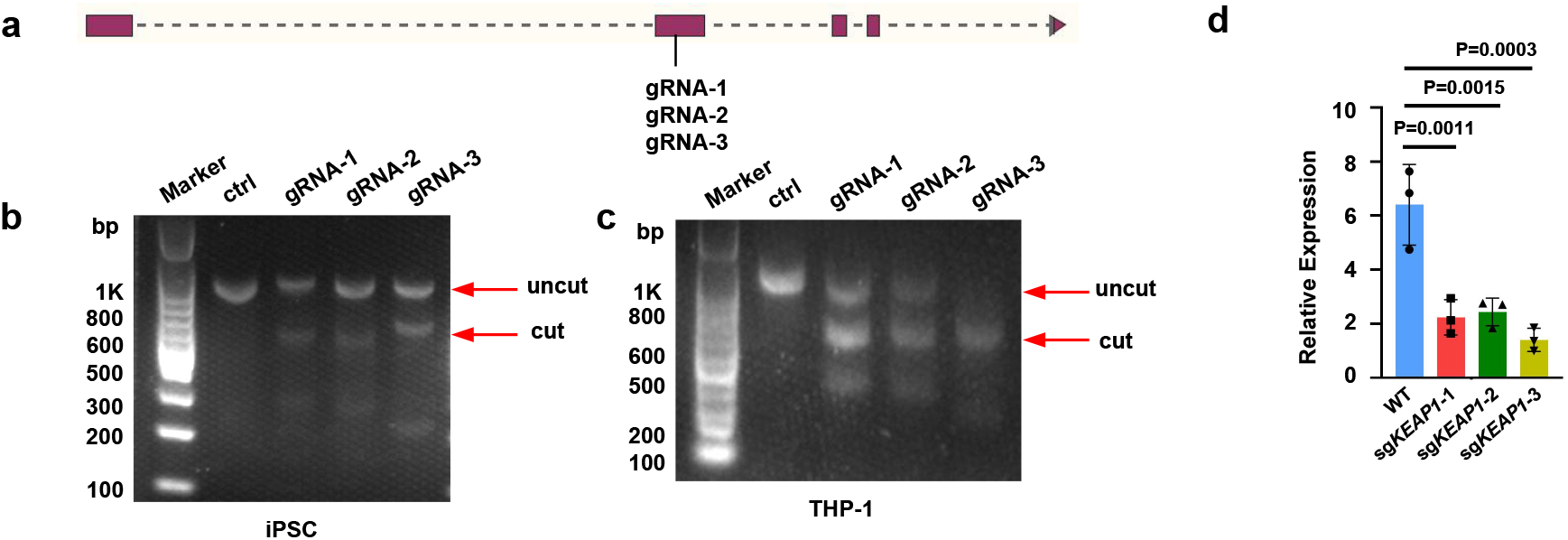
*KEAP1* deletion in THP-1 cells, related to **Fig. 1**. **a**, CRISPR-Cas9-mediated *KEAP1* KO using three sgRNAs targeting exon 2 of the *KEAP1* gene. **b,c**, Validation of DNA cleavage efficiency by T7 endonuclease assays in iPSCs (b) and THP-1 cells (c). **d**, Relative expression of *KEAP1* in WT and sg*KEAP1* transfected tMACs (n=3). Data was shown as mean ± SD. Statistics by one-way ANOVA test.

**Extended Data Fig. 3.**
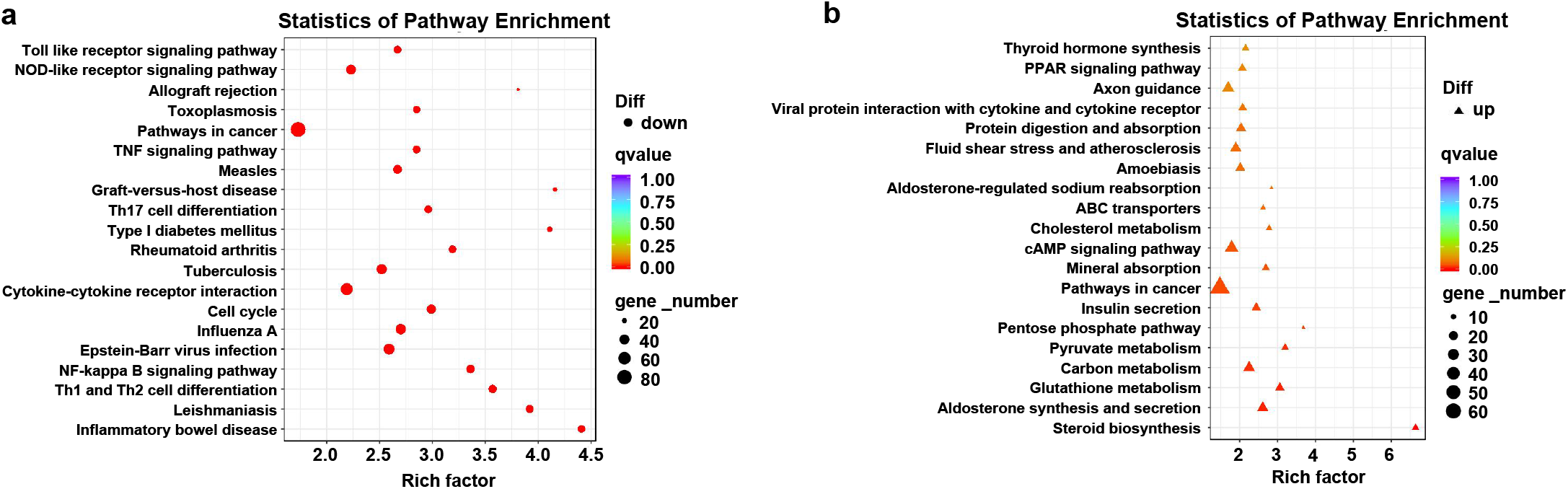
Pathway enrichment in *KEAP1*-deleted macrophages, related to **Fig. 1**. **a,b**, Top enriched gene sets down-regulated (**a**) or up-regulated (**b**) in sg*KEAP1*-3-transduced tMACs compared to sgControl-transduced cells after LPS and IFN-γ stimulation for 8 h.

**Extended Data Fig. 4.**
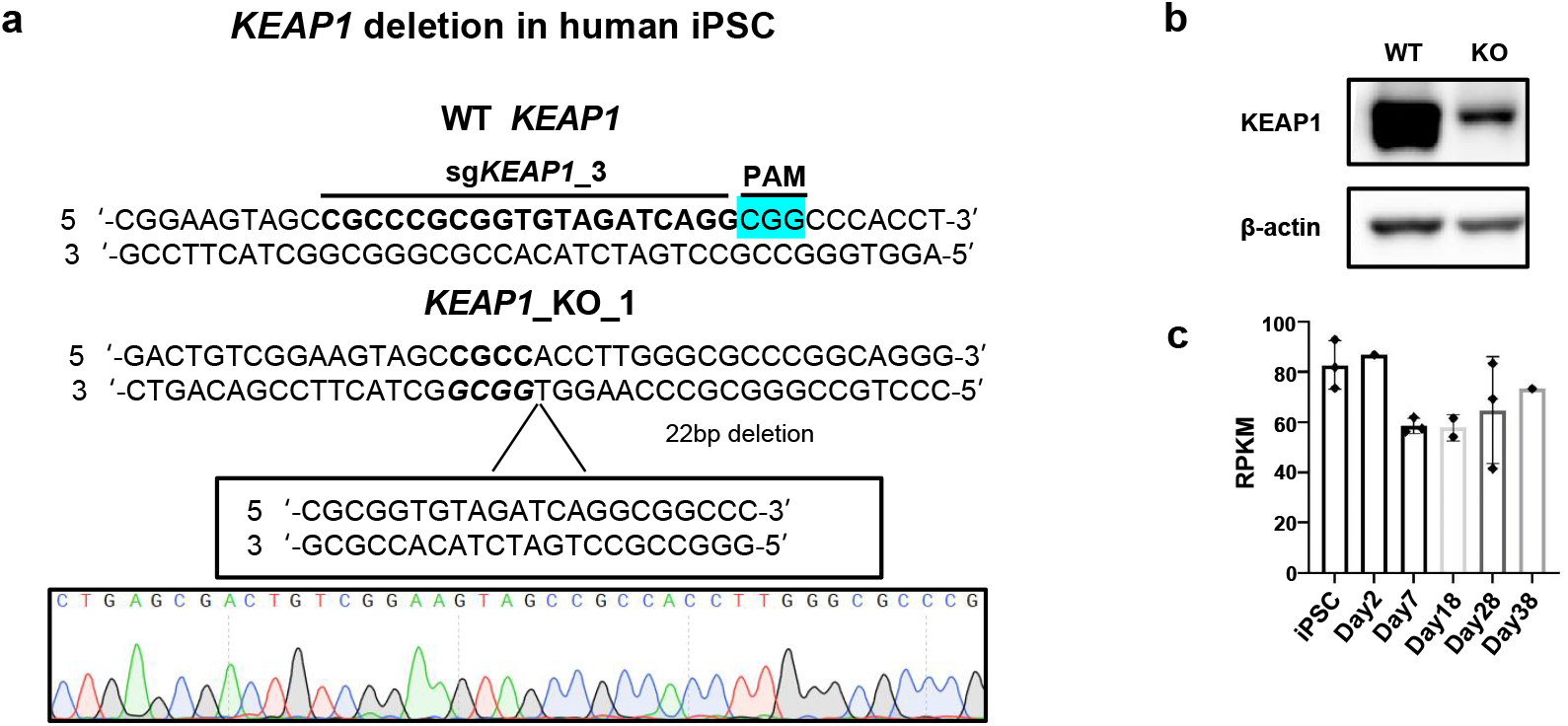
*KEAP1* deletion in human iPSCs, related to **Fig. 1**. **a**, Comparison of the DNA sequence in the *KEAP1* KO iPSC clone (by Sanger sequencing) with *KEAP1* WT sequence, revealing a 22 bp deletion in the gRNA targeted region. **b**, The protein expression of *KEAP1* in WT and *KEAP1* KO iPSCs was evaluated by western blotting. **c**, RNA-seq data for the expression of *KEAP1* in iPSCs and differentiated cells on day 2, 7, 18, 28, and 38 (iPSC, n=3; Day 2, n=1; Day 7, n=3; Day 18, n=2; Day 28, n=3; Day 38, n=1). Data was shown as mean ± SD.

**Extended Data Fig. 5.**
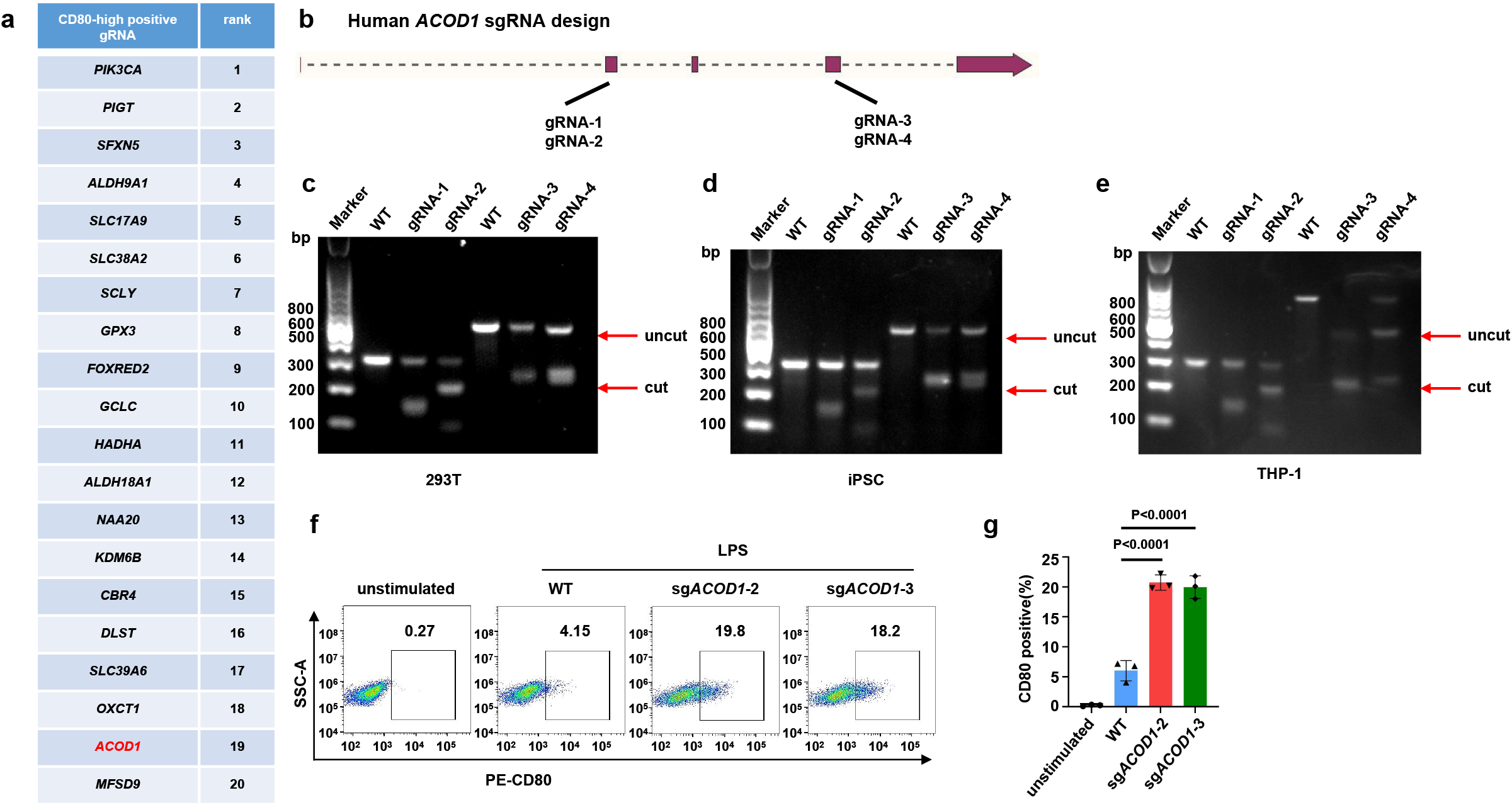
*ACOD1* deletion in human macrophages resulted in enhanced pro-inflammation activation, related to **Fig. 2**. **a**, Top 20 sgRNA targeted genes enriched in the CD80-high population identified by a CRISPR Screen in iPSC-derived macrophages. **b-e**, CRISPR/Cas9-mediated *ACOD1* knockout using four sgRNAs located in exons 2 and 4 of the *ACOD1* gene, and validation of DNA cleavage efficiency by T7 endonuclease assays in 293T (**c**), iPSC (**d**), and THP-1 cells (**e**). **f,g**, Flow cytometry plots and quantification of CD80 expression in unstimulated, WT, and sg*ACOD1* transduced tMACs with 50 ng/mL LPS stimulation for 24 h (g, n=3). Data was shown as mean ± SD. Statistics by one-way ANOVA test.

**Extended Data Fig. 6.**
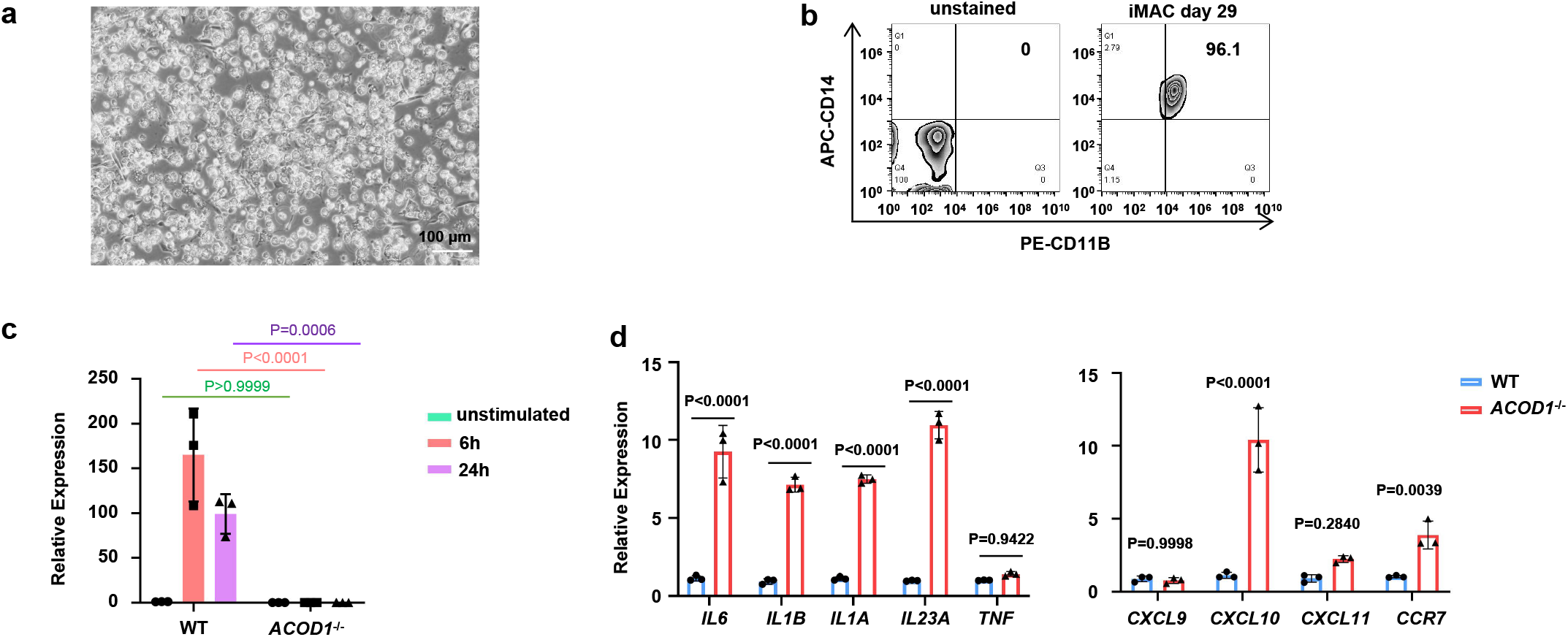
*ACOD1* deletion resulted in elevated pro-inflammatory gene expression in iMACs, related to **Fig. 3**. **a**, Representative images of differentiated iMACs at day 29. **b**, CD14 and CD11B expression on iMACs at day 29 was determined by flow cytometry. **c**, The relative expression of *ACOD1* in WT and *ACOD1*^-/-^ iMACs with the indicated treatments, including unstimulated, and 50 ng/mL LPS plus 50 ng/mL IFN-γ stimulation for 6 and 24 h (n=3). **d**, qRT-PCR for mRNA expression of pro-inflammatory genes and anti-inflammatory genes in WT and *ACOD1*^-/-^ iMACs after LPS and IFN-γ stimulation for 24 h (n=3). **c,d**, Data was shown as mean ± SD. Statistics by two-way ANOVA test.

**Extended Data Fig. 7.**
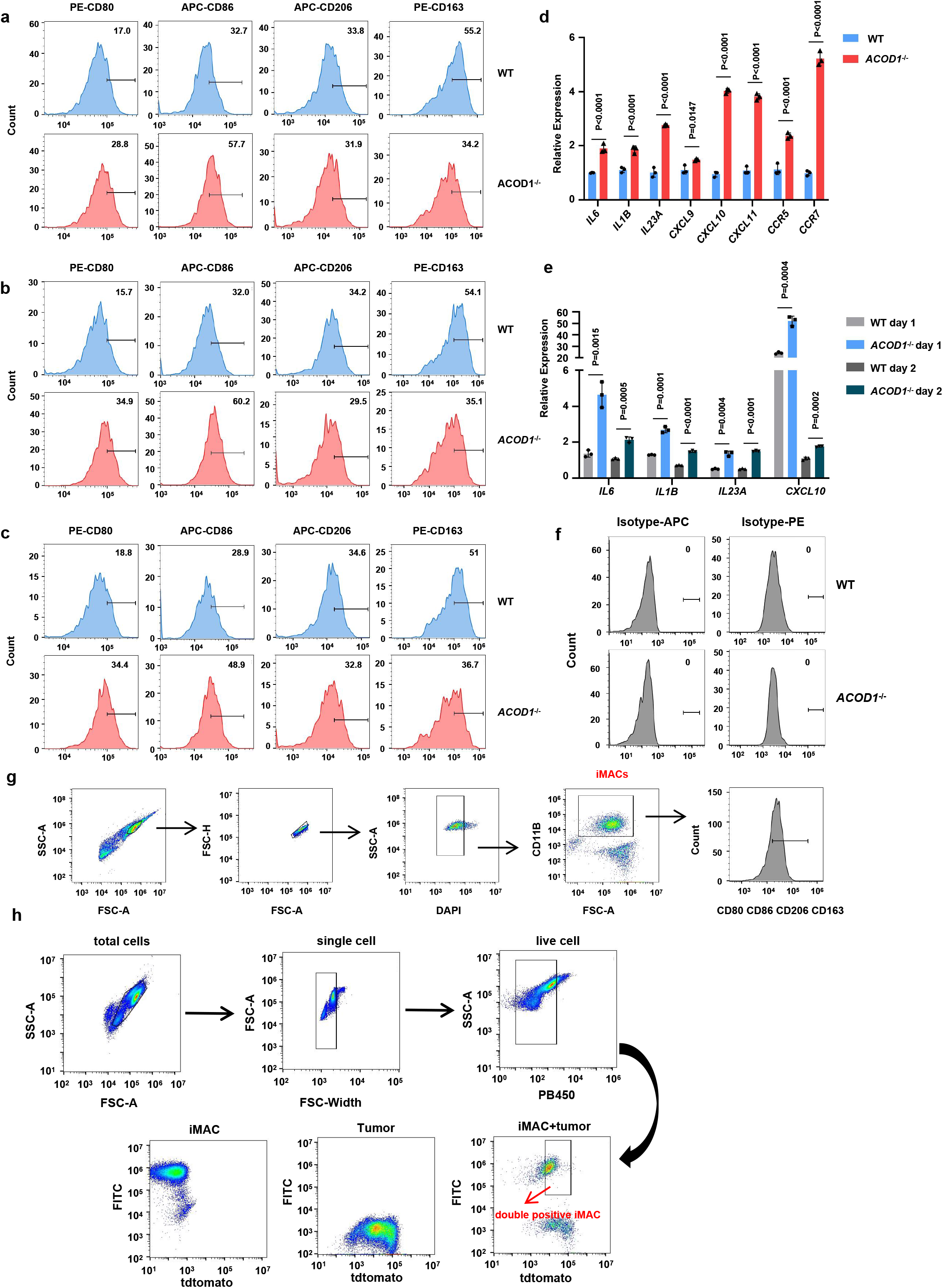
*ACOD1*^-/-^ iMACs exhibited increased pro-inflammatory activation when co-cultured with tumor cells, related to **Fig. 4**. **a-c**, The expressions of CD80, CD86, CD163, and CD206 in WT or *ACOD1*^-/-^ iMACs after co-cultured with (**a**) Nalm6 (E:T=3:1), (**b**) K562 (E:T=5:1) or (**c**) K562 (E:T=3:1) for 24 h measured by flow cytometry and displayed as histograms. **d**, qRT-PCR for mRNA expression of pro-inflammatory genes in WT and *ACOD1*^-/-^ iMACs after co-culture with Nalm6 (E:T=3:1) for 24 h (n=3). Data was shown as mean ± SD. **e**, qRT-PCR for mRNA expression of pro-inflammatory genes in WT and *ACOD1*^-/-^ iMACs after co-culture with Nalm6 (E:T=5:1) for 24 h (day 1) or 48 h (day 2) (n=3). Data was shown as mean ± SD. Statistics by two-way ANOVA test. **f**. WT or *ACOD1*^-/-^ iMACs were stained by APC or PE isotype and displayed as histograms. **g**, Gating strategy of CD80-high, CD86-high, CD163-high, or CD206-high cells. **h**, Gating strategy of the phagocytosis assay. The iMAC cells were stained with a green dye and thus they were positive in the green channel, and the tumor cells were transduced with tdtomato, and thus there were positive in the red channel. The iMAC cells undergoing phagocytosis were those showing double positive, compared with the single positive iMAC cells or tumor cells.

**Extended Data Fig. 8.**
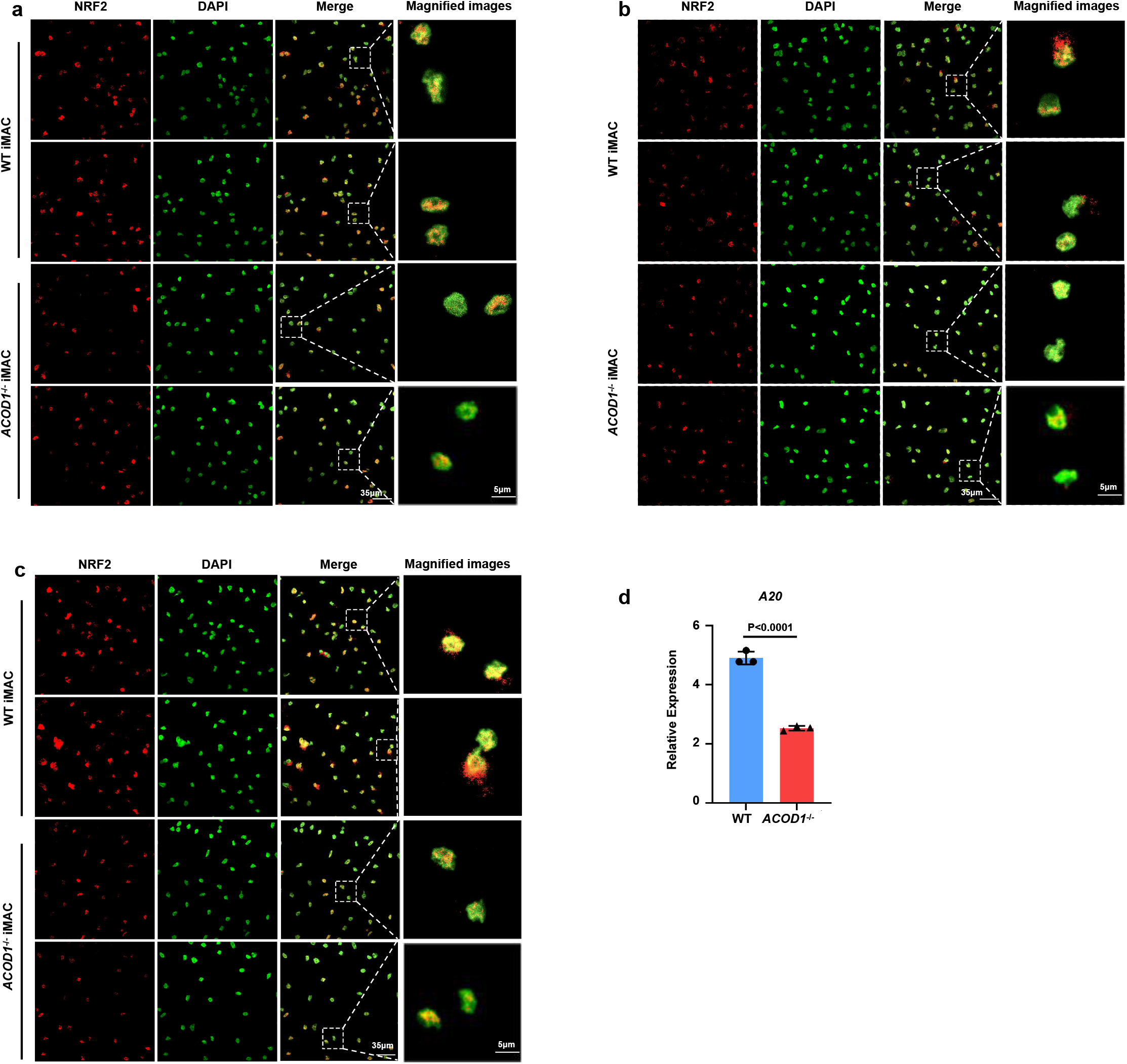
*ACOD1* deletion resulted in decreased nuclear expression of NRF2 and decreased expression of the NF-κB pathway negative regulator *TNFAIP3* (*A20*), related to **Fig. 5**. **a-c**, Representative confocal images of NRF2 in WT and *ACOD1*^-/-^ iMACs after LPS and IFN-γ stimulation for (**a**) 0 h, (**b**) 30 min, or (**c**) 8 h. **d**, qRT-PCR for mRNA expression of *TNFAIP3* (*A20*) in WT and *ACOD1*^-/-^ iMACs after LPS and IFN-γ stimulation for 24 h (n=3). Data was shown as mean ± SD. Statistics by unpaired t test.

**Extended Data Fig. 9.**
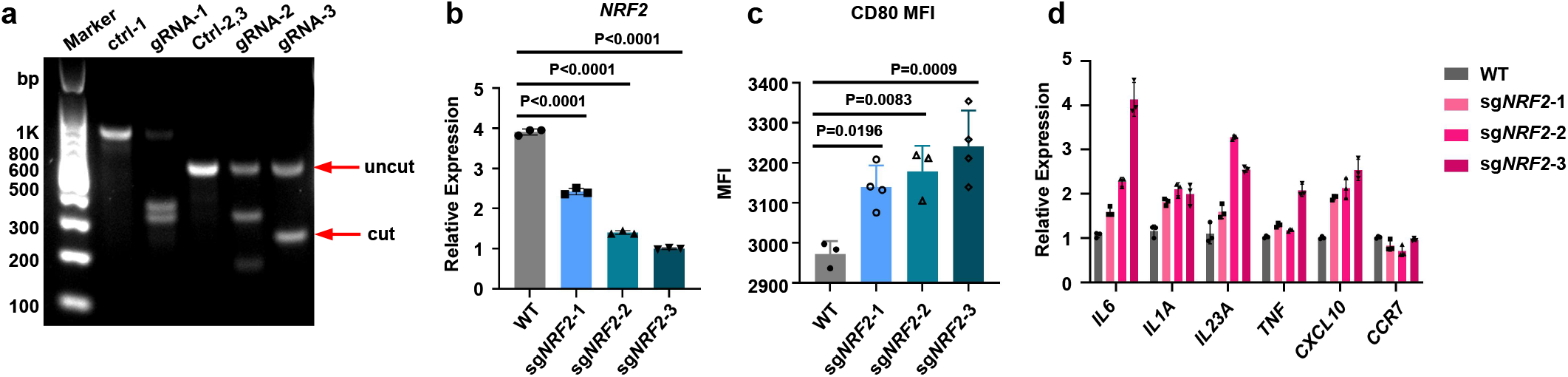
*NRF2* deletion promoted pro-inflammatory activation in tMACs, related to **Fig. 5**. **a**, Validation of DNA cleavage efficiency by T7 endonuclease assays in THP-1 cells. **b**, qRT-PCR for mRNA expression of *NRF2* in WT and sg*NRF2*s transduced THP-1 cells (n=3). **c**, Quantification of CD80 MFI measured by flow cytometry in WT and sgNRF2s transduced macrophages after LPS and IFN-γ stimulation for 24 h (WT, n=3; sg*NRF2*-1, n=4; sg*NRF2*-2, n=3; sg*NRF2*-3, n=4). **d**, qRT-PCR for mRNA expression of pro-inflammatory genes in WT and sg*NRF2*s transduced macrophages after LPS and IFN-γ stimulation for 24 h (n=3). **b-d**, Data was shown as mean ± SD. **b,c**, Statistics by one-way ANOVA test.

**Extended Data Fig. 10.**
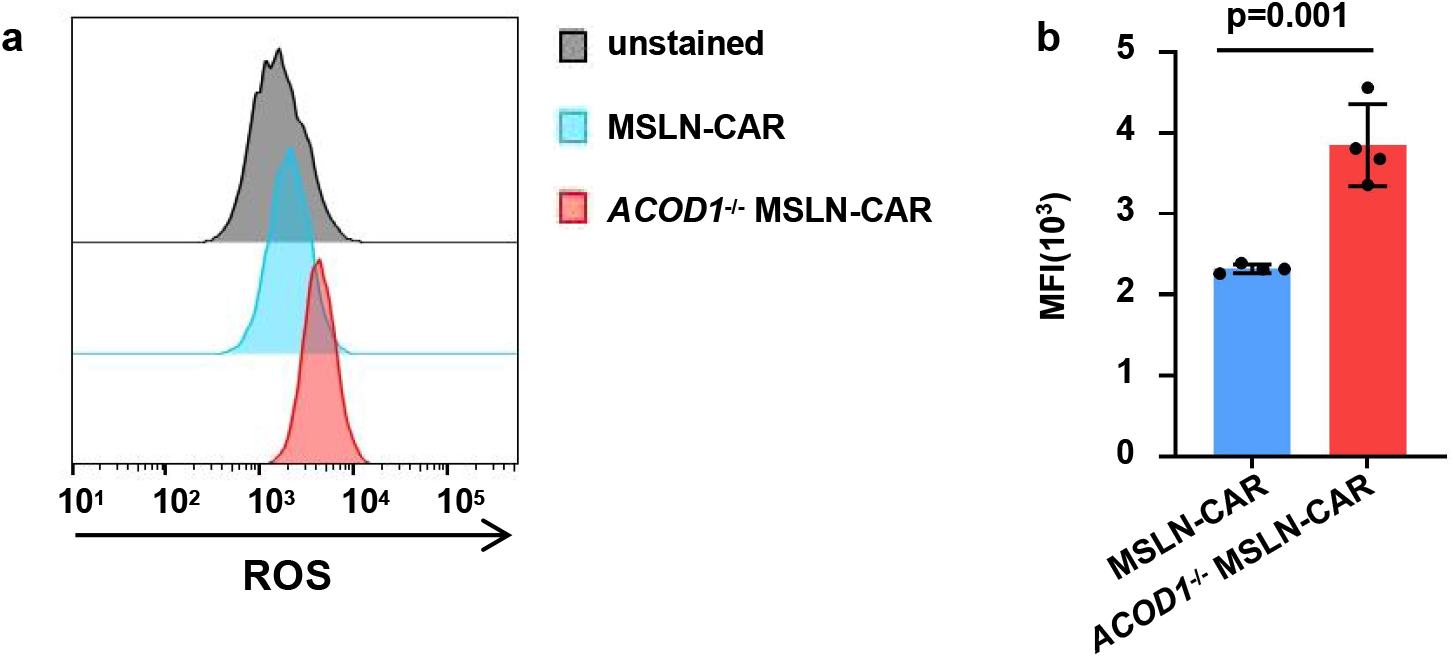
ACOD1 deletion promoted ROS production in iMACs, related to Fig. 6. **a,b,**ROS in MSLN-CAR and *ACOD1*^-/-^ MSLN-CAR-iMACs (a) and mean fluorescence intensity (MFI) quantification (b) was determined by flow cytometry after stimulated by LPS plus IFN-γ (50 ng/mL each) for 24 h which were stained by MitoSOX Red Mitochondrial Superoxide Indicator. **b**, Data were shown as mean ± SD (n=4), Statistics by unpaired t test.

**Extended Data Fig. 11.**
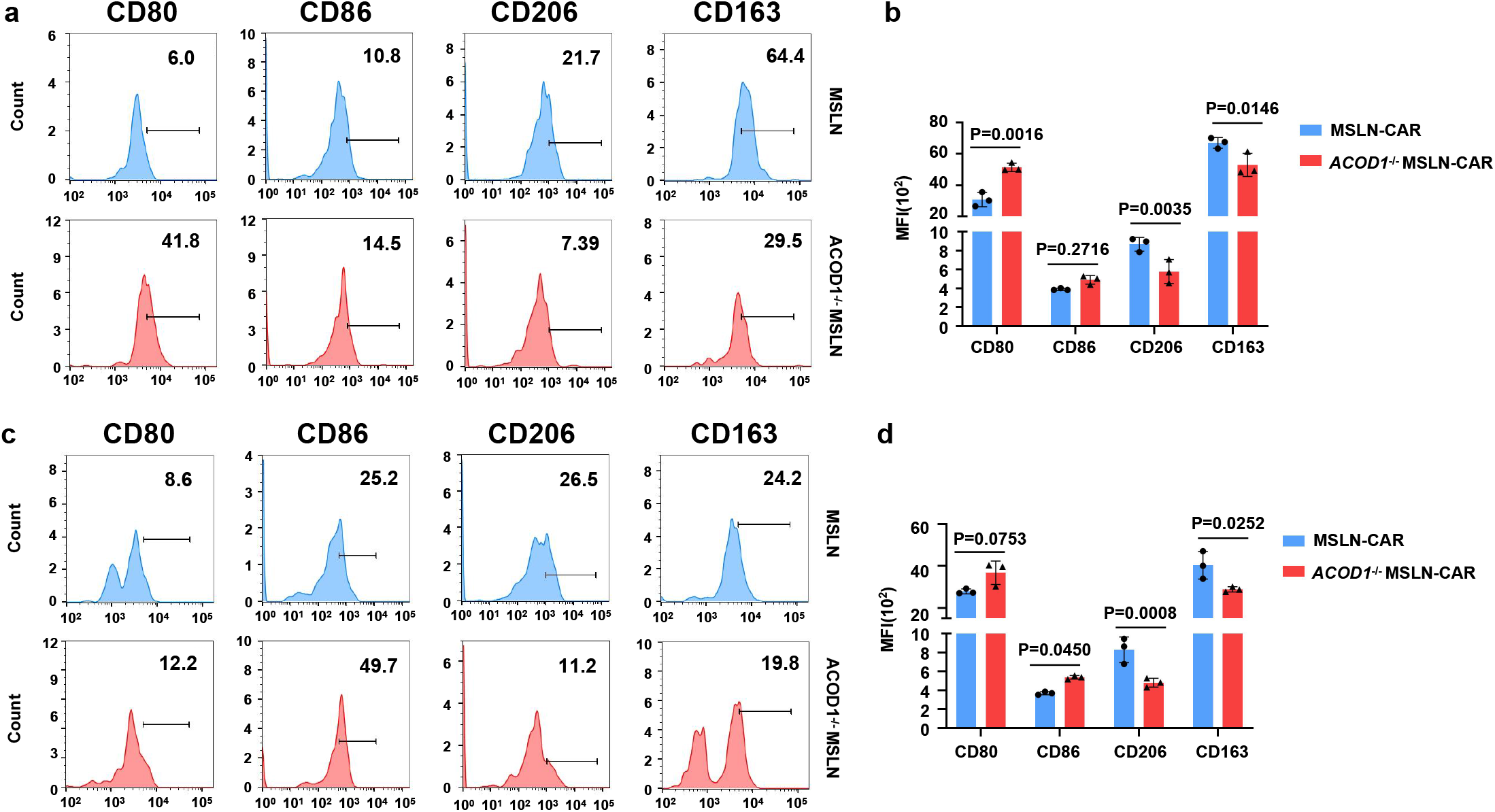
*ACOD1* deletion promoted pro-inflammatory activity of iMACs *in vivo*, related to Fig. 6. **a,b**, Subcutaneous tumor model was established in NSG mice. 7 days later, MSLN or *ACOD1*^-/-^ MSLN-CAR-iMACs were injected intratumorally. After 7 days, the expression of CD80, CD86, CD206, and CD163 in MSLN or *ACOD1*^-/-^ MSLN-CAR-iMACs was measured by flow cytometry and the representative data was displayed as histograms (**a**). Data averaged from three independent experiments were shown as mean ± SD (**b**) (n=3). **c,d**, After 14 days, the expression of CD80, CD86, CD206, and CD163 in MSLN or *ACOD1*^-/-^ MSLN-CAR-iMACs was measured by flow cytometry and the representative data was displayed as histograms (**c**). Data averaged from three independent experiments were shown as mean ± SD (**d**) (n=3). **b,d**, Statistics by two-way ANOVA test.

**Extended Data Fig. 12.**
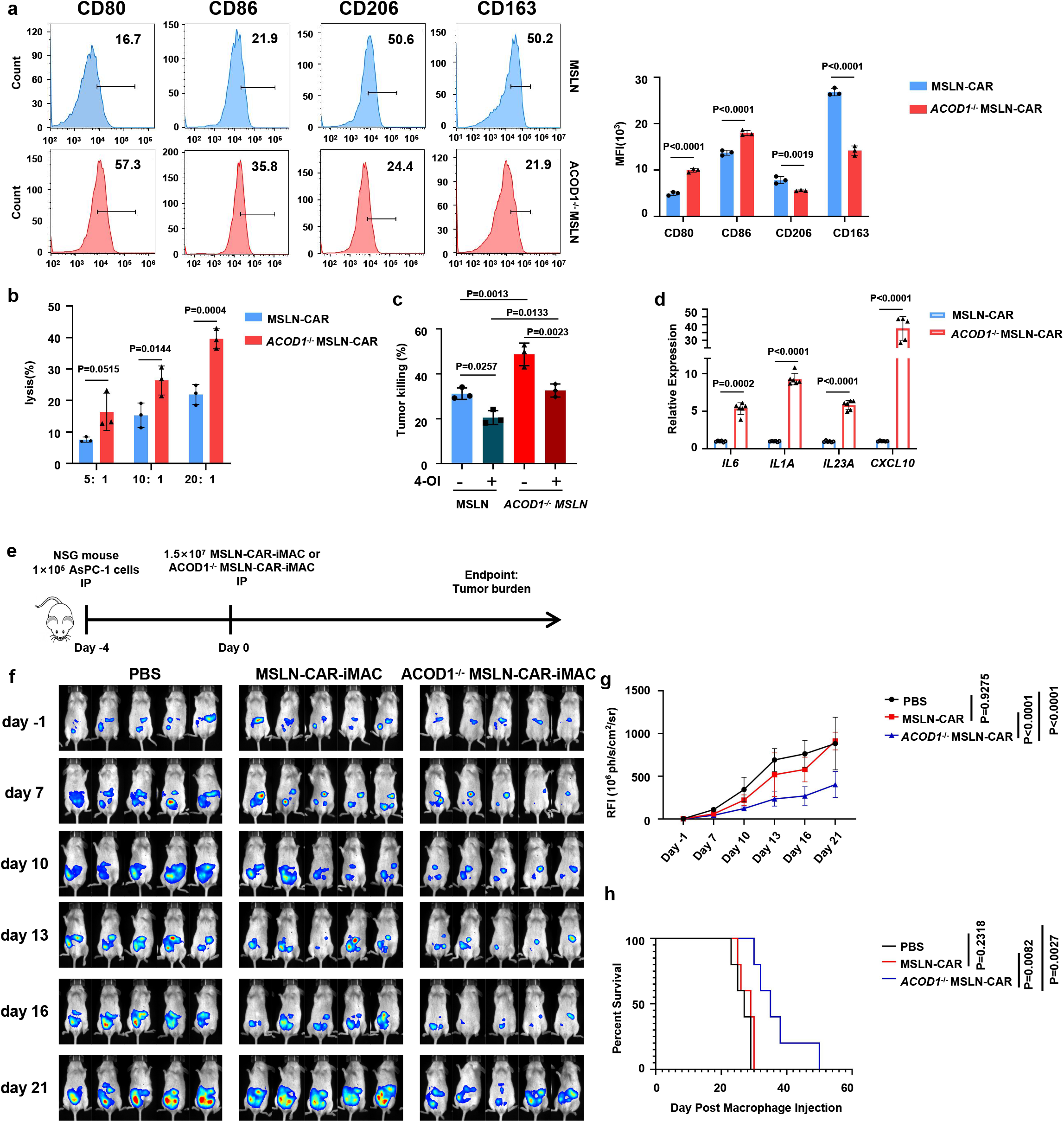
*ACOD1* deletion promoted anti-pancreatic cancer activity of iMACs *in vitro* and *in vivo*, related to **Fig. 6**. **a**, The expression of CD80, CD86, CD206, and CD163 in MSLN or *ACOD1*^-/-^ MSLN-CAR-iMACs after co-cultured with pancreatic cancer cell AsPC-1 (E: T=5:1) for 24 h measured by flow cytometry and the representative data was displayed as histograms (left). Data averaged from three independent experiments were shown (right) (n=3). **b**, Luciferase assays for CAR-iMAC cytotoxicity activity against cancer cells when co-cultured with AsPC-1 cells for 24 h (E: T=5:1, 10:1, or 20:1) (n=3). **c**. Luciferase assays for CAR-iMAC cytotoxicity activity against cancer cells with or without 4-OI (250 μM) when co-cultured with AsPC-1 cells for 24 h (E: T=10:1) (n=3). **d**, qRT-PCR for mRNA expression of pro-inflammatory genes in MSLN or *ACOD1*^-/-^ MSLN-CAR-iMACs after co-cultured with AsPC-1 cells (E: T=5:1) for 24 h (n=3). (a-c) Data was shown as mean ± SD. Statistics by two-way ANOVA test. **e**, A diagram of the in vivo treatment scheme. **f**, IVIS images showing progression of tumor (n=5 per group). **g**, Tumor burden on day -1, 7, 10, 13, 16 and 21 was quantified and displayed as mean ± SD. Statistics by two-way ANOVA test. **h**, The Kaplan-Meier curve demonstrating survival of the mice. Statistics by two-tailed log-rank test.

**Extended Data Fig. 13.**
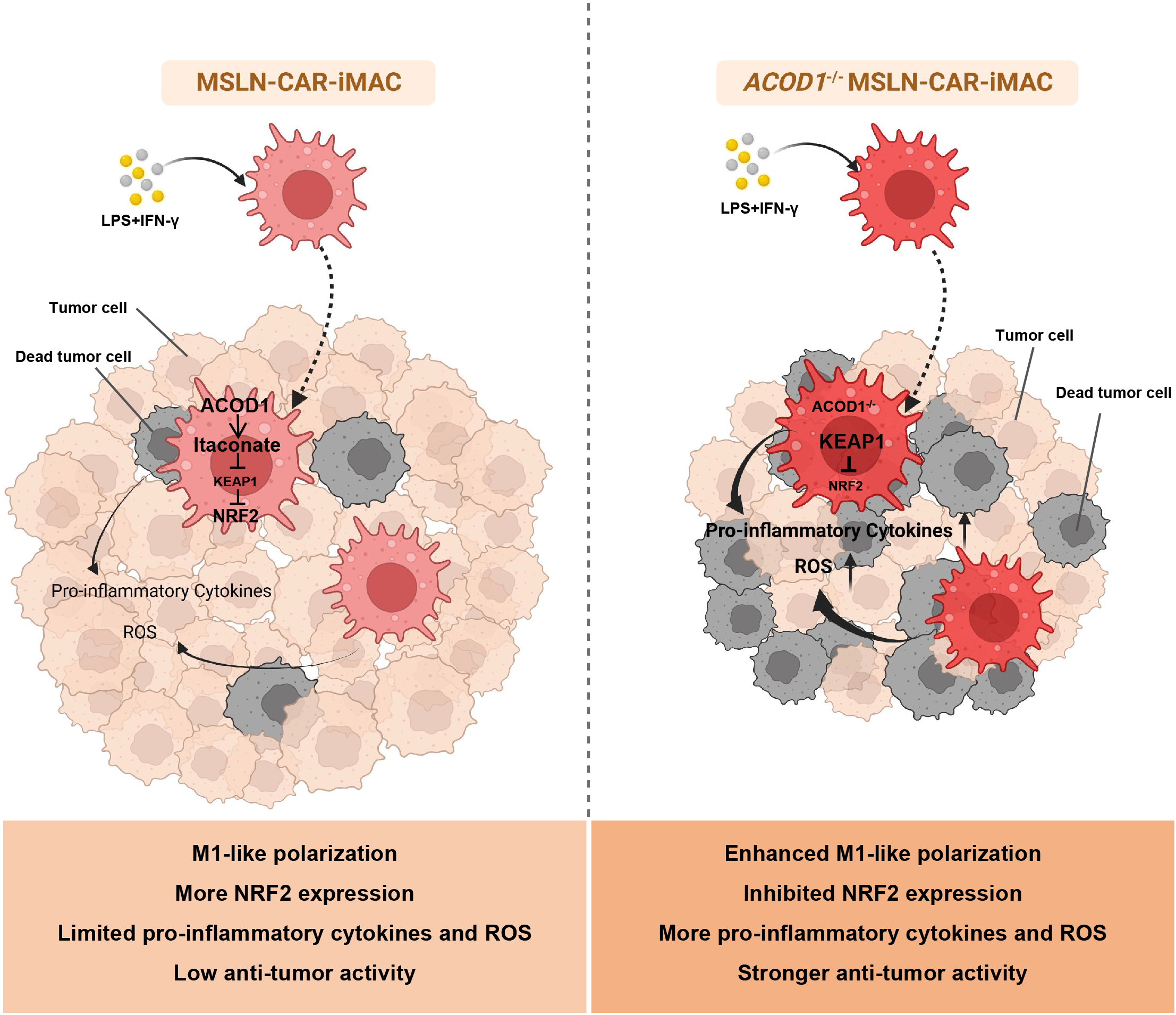
The diagram of ACOD1 regulating the anti-tumor effect of MSLN-CAR-iMACs, MSLN-CAR-iMACs and *ACOD1*^-/-^ MSLN-CAR-iMACs were activated upon stimulation with LPS and IFN-γ. The expression of itaconate was abrogated by *ACOD1* deletion in *ACOD1*^-/-^ MSLN-CAR-iMACs. Itaconate is known to alkylate cysteine residues on KEAP1, promoting the accumulation and nuclear translocation of NRF2, which leads to the expression of downstream genes with anti-oxidant and anti-inflammatory properties. Consequently, *ACOD1*^-/-^ MSLN-CAR-iMACs showed lower expression of NRF2, but higher levels of pro-inflammatory cytokines and ROS. Furthermore, these cells exhibited enhanced M1-like polarization and stronger anti-tumor activity.

